# Electrophysiological development and functional plasticity in dissociated human cerebral organoids across multiple cell lines

**DOI:** 10.1101/2025.03.24.644896

**Authors:** Adam Pavlinek, Lucia Dutan Polit, Roland Nagy, APEX consortium, Madeline A. Lancaster, Anthony C. Vernon, Deepak P. Srivastava

**Author notes:** Co-Senior authors (Anthony C. Vernon, Deepak P. Srivastava).

## Abstract

Microelectrode arrays (MEAs) are increasingly used to profile the development of synchronised activity in neural organoids, yet no organoid study has investigated the consistency of electrophysiological development across cell lines. Here, we used dissociated neural organoids derived from four cell lines on MEAs to characterise functional synapse development using multiple parameters across time. The dissociated organoids had increasing functional connectivity and network activity over time across all cell lines and plasticity in response to synaptic-like stimulation. Like the organoids they were derived from, dissociated organoid cultures contained a diverse mixture of cell types. Variability in activity parameters was associated with differences in cell type composition and regional identity, which in turn were affected by donor cell line and batch effects. These results demonstrate that dissociated cerebral organoids can generate functional neurons, akin to primary neuronal cultures from brain tissue, providing a scalable model for studies of neurodevelopment and synaptic function.

## INTRODUCTION

Much of what is known about electrical activity in the developing human brain is inferred from rodent models, limiting our understanding of species-specific mechanisms that shape neural circuit function.^1^ Despite evolutionary conservation^2–4^, the human cortex exhibits unique features distinguishing it from other primates and mammals. This includes differences in cytoarchitecture, greater neuron density and diversity^5,6^, cell type differences^5,7,8^, and increased connectivity^9,10^. In the human prefrontal cortex, human pyramidal neurons have more synapses and dendritic spines, greater spine volume and neck length as compared to mouse and primate pyramidal neurons^10^. These differences contribute to altered information processing and electrophysiological properties, including prolonged synaptic maturation and distinct patterns of neuronal excitability.^3,9,10^ Many of these species differences originate in neurodevelopment.^10–12^ Synapse development and maturation are protracted in humans^13^, with human neurons showing neoteny, even when transplanted into mice^14–16^. Neoteny appears to be cell-intrinsic^10^ and driven by human-specific genetic and epigenetic differences^17,18^. Recent advances in human-derived cellular models such as human induced pluripotent stem cell (iPSC) -derived neurons and neural organoids offer an opportunity to study the development and complexity of human cortical development, particularly the development of human neural networks and their emergent network properties.^19–21^

Micro-electrode arrays (MEAs) have widely been used to investigate the development of 2 dimensional (2D) human neural networks and their plasticity.^20–23^ In monocultures of iPSC neurons, initially random spiking and bursting can gradually become more complex^20^, with emergence of properties such as network bursting, and oscillations that are indicative of synaptic connectivity.^22,24–26^ Such spontaneous neuronal activity is thought to aid the development of synchronised neuronal activity, refinement of synaptic connections and network maturation.^27,28^ As well as investigating normative development, MEA recordings can be used to study disease-specific phenotypes^20^ including emergent phenotypes using iPSC models of neurodevelopmental conditions, such as autism^29–31^. However, while 2D monocultures are amenable to electrophysiological analysis, they lack the cellular diversity of the developing human brain, with some studies utilising rodent astrocytes to enhance electrophysiological development^20,21^.

Three-dimensional (3D) human organoid models offer a more physiologically relevant platform for studying human-specific neurodevelopment.^32^ Specifically, transcriptomic analyses across different differentiation methods have shown that major gene expression programmes, cell types, and epigenetic regulation in the developing brain are better preserved in organoids when compared to fetal tissue, than in 2D-cultures^33–38^, at least up until the late second trimester of fetal development^39^. This includes human-specific developmental changes, such as delayed neuronal maturation and human-specific gene expression^8^. Different protocols now exist for the generation of neural organoids, which can be categorised as either unguided (relying on intrinsic patterning of tissue) or regionalised (guided by exogenously added factors).^35,40^ Whilst unguided neural organoids have increased variability between organoids as compared to more guided approaches, they also have greater cellular diversity.^41^

Human neural organoids display spontaneous electrical activity, and develop synchronised network activity, measurable using calcium imaging, patch-clamp recordings^42,43^, and MEAs^44–47^. These studies have typically characterised the neuronal networks in organoids and their dynamics over time in limited numbers of organoids from N=1 or 2 donor cell lines ^43–45,48^. Fair *et al.* ^45^ observed increasing rates of spiking in cerebral organoids from one cell line with increasing developmental time up to 161 days in culture, which correlated with the appearance of greater neuronal and astrocyte diversity^45^. Similarly, Giandomenico *et al.* ^46^ observed network bursting in organoid slices from the H9 embryonic stem cell (ESC) line. Despite these advances, the electrophysiological properties of neural organoids remain incompletely characterized, particularly in scalable, high-throughput formats. Addressing this gap requires the adaptation of organoid-derived neuronal cultures for scalable MEA-based studies to systematically assess their network activity, connectivity, and responsiveness to physiological stimuli across multiple cell lines and developmental timepoints. No study has systematically studied the effects of organoid batch, cell line, and chromosomal sex on the development of electrophysiological activity, as measured by MEAs, in unguided organoid models. Understanding batch and donor line variability in MEA readouts is essential for informing the design and interpretation of studies that use patient-derived or genetically engineered lines. In addition, whilst there is evidence that organoid electrical activity changes in response to drugs, such as tetrodotoxin ^46^ or diazepam ^49^, the effects of more physiological stimuli, such as those for induction of long-term potentiation^50^ or steroid hormones have not been investigated.

In this study, we use organoid dissociation and 2D culture as a scalable approach for longitudinal recording of activity over longer time periods. Unlike neuronal monocultures, dissociated organoid cultures benefit from the increased cell type diversity of organoids and undergo initial patterning and development in a 3D context. We systematically characterise the development of spontaneous electrophysiological activity and network properties in dissociated unguided cerebral organoids derived from four donor cell lines. By leveraging MEA recordings, we assess how batch effects, donor cell line, and chromosomal sex influence network maturation over time. We further evaluate the responsiveness of dissociated organoid-derived networks to chemical long-term potentiation (chLTP) induction^50^ and the steroid hormone 17β-estradiol (E2), a key modulator of synaptic plasticity in rodent models^51–57^. We show that despite batch-dependent variability, dissociated organoid cultures exhibit consistent temporal progression in network properties, supporting their use as a scalable and reproducible model for studying human neural network activity. Moreover, by investigating how organoid-derived neuronal networks respond to plasticity-inducing stimuli and hormonal modulation, this study provides a critical foundation for future research into the mechanisms underlying human-specific synaptic development and function.

## RESULTS

### Dissociation of cerebral organoids and culture on MEAs

To analyze the development of electrical activity over time in neural cultures from dissociated organoids, we generated unguided neural organoids from two ESC and two iPSC cell lines and cultured them up to 45 days *in vitro* (DIV) using the established “cerebral organoid” method ^46,58^, originally developed by Lancaster et al. ^59^ (Figure 1A). We used organoids with visible cortical plate regions (Figure 1B) from batches that contained organoids with dorsal forebrain tissue (Figure 1C), using established characterisation markers^59–61^. At 40 DIV, cerebral organoids have early-born MAP2-positive neurons on the outside of neuroepithelial regions, which in dorsal forebrain regions were positive for TBR1 and CTIP2 (Figure 1D). Random spiking, with physiological waveforms, was detected in a subset of whole 40 DIV organoids attached to an MEA (Figure S1) but with no detectable network activity. To reduce inter- organoid variability, we pooled multiple organoids per batch together (a minimum of 6 organoids), dissociated them into single cells using a papain-based dissociation approach (Figure 1A), and then plated them onto multi-well plates with 16 embedded microelectrodes per well (Figure 1E). Across dissociated cultures from the four cell lines, we observed that cells were initially evenly distributed in the well but became more aggregated over time (Figure 1F). At 13 days post dissociation (DPD), the cells had visible processes but also had some early signs of uneven aggregation, with denser areas of cells in the well. By 27 DPD, there was extensive clustering of cells, with bundles of axons extending between clusters, the clusters then increased in size between 27 and 37 DPD. The dissociated organoid cultures developed spontaneous activity and clustered bursting within 20 DPD (Figure 1G, 1H). We used established analysis pipelines^26^ to extract parameters (Table S1) to analyse the characteristics of activity and connectivity in the dissociated cultures (Figure 1I).

**Figure 1.**
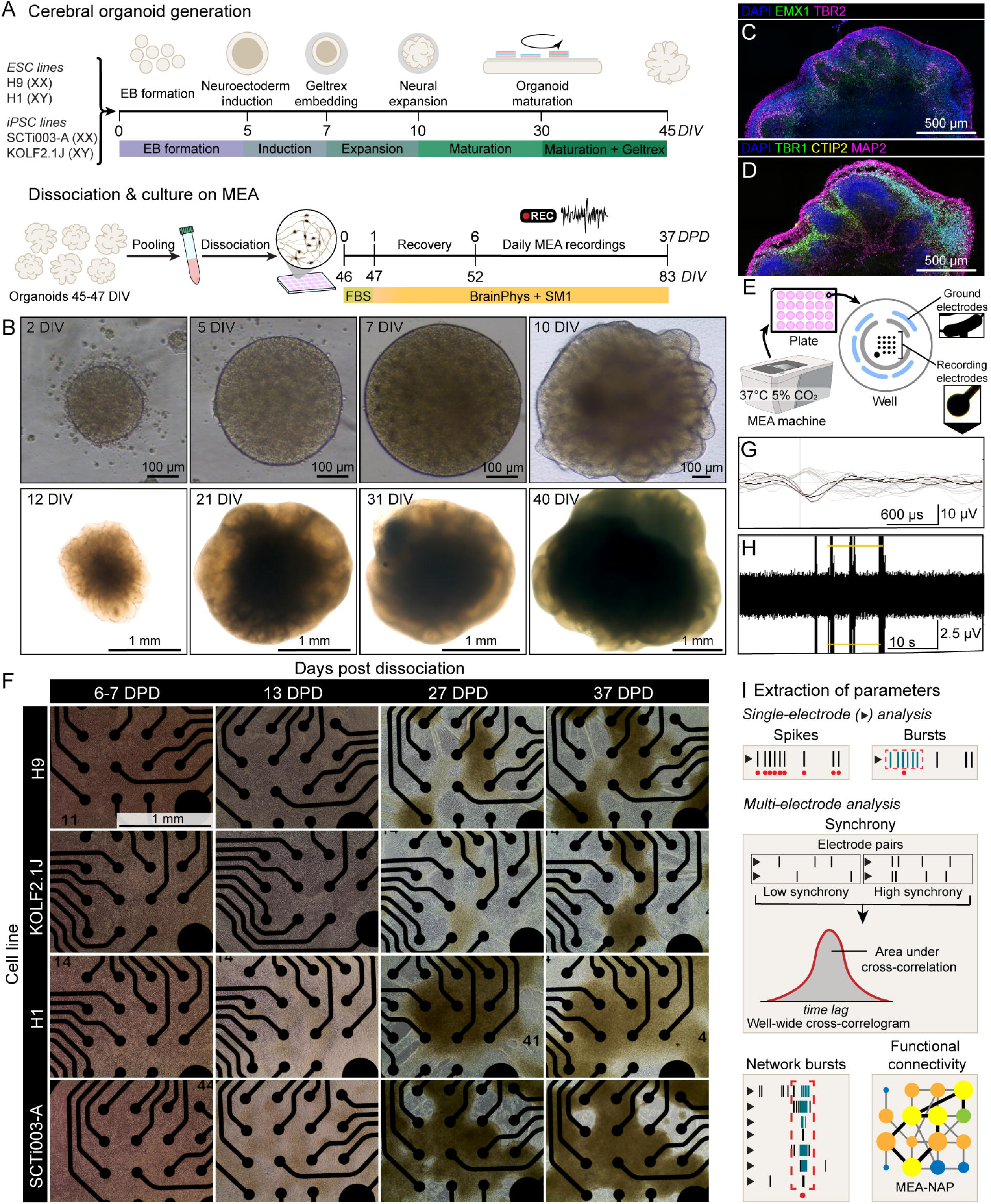
Overview of dissociated organoid culture, morphology, and MEA recording. (A) Organoid generation and dissociated culture timeline. Top: generation of cerebral organoids with days *in vitro* (DIV). Bottom: organoid dissociation and culture. Timing from the day of dissociation shown as days post dissociation (DPD) The colored bar below the timelines indicates the culture medium used (see methods for details). (B) Brightfield images of organoid development hallmarks. Images from the embryoid body (EB) stage (0-5 DIV), neural induction (7 DIV), expansion of neuroepithelial buds following Geltrex embedding (10 DIV), and organoid maturation (12 DIV - 40 DIV). At 31 DIV, extended neuroepithelial regions can be seen at the edges. Images show different representative H9 line organoids. (C-D) Composite images of neural rosettes from confocal tile scans of two adjacent serial sections of a representative SCTi003-A line organoid at 40 DIV. Immunostaining for dorsal forebrain markers EMX1 and TBR2 (D), MAP2, CTIP2 and TBR1 (E). Brightness and contrast adjusted for clarity. See also Figure S1. (E) Schematic of recording set-up with 24-well MEA plate showing within-well arrangement of ground and recording electrodes. (F) Representative brightfield images of dissociated organoids on 16-electrode MEA. Each column shows 6-7-, 13-, 27-, and 37-DPD. Each row shows images from one batch of each cell line. All images are at the same scale. (G) Snapshot of recording from H9 line dissociated organoids at 34 DPD (80 DIV) showing unsorted overlaid spike waveforms recorded at a single electrode. Waveforms correspond to individual spikes above the 5.5 standard deviation (SD) threshold. The darkest waveform represents the most recent spike recorded at the electrode. (H) Representative time segment at 34 DPD showing raw neural spikes recorded at one electrode, in the same well and time as in (G). Horizontal lines indicate the 5.5 SD threshold. (I) Schematic summary of key single electrode and multi-electrode neuronal activity parameters. Parameter details in Table S1. Abbreviations: MEA-NAP, MEA network analysis pipeline^26^

### Dissociated organoids from four donor cell lines develop network connectivity over time

To determine the dynamics of electrophysiological activity over time in the dissociated organoid cultures, we analysed changes in activity between 6 DPD when we could detect spiking activity, and 37 DPD across six core parameters (Table S1). We generally observed increases in firing rate and bursting over time (Figure 2A), however this varied between batches. For instance, mean firing rate was increased at 37 DPD compared to 6 DPD in both H9 and KOLF2.1J line dissociated organoids but we did not see a significant increase in the H1 or SCTi003-A lines (Figure 2A). We saw more consistent increases in connectivity over time, including increasing area under normalized cross-correlation (AUNCC), as well as increasing node degree. Both node degree and AUNCC were higher at 37 DPD compared to 6 DPD across all lines (Figure 2B, 2C, Table S3). Correspondingly, regular network bursts occurred in dissociated organoids from all cell lines (Figure 2D). Overall, we saw increasing number and strength of connections over time and greater participation of nodes in the network (Figure 2E). However, there was variability between wells in these parameters (Figure 2F-H). For example, only a subset of wells developed network bursting in H1 and SCTi003-A dissociated cultures (Figure 2H). There were cell line and batch differences; cells from the KOLF2.1J line developed the highest AUNCC and greatest number of network bursts, followed by the H9 line (Figure 2G-H). At 37 DPD, there were no significant differences between XX and XY lines in any of the six core parameters (Table S3). We did see greater node degree (F1,100=6.07, p=0.0155, Table S3) in ESC networks (10.65±3.36 mean connections) compared to iPSC networks (7.38±4.33 mean connections) but no iPSC/ESC line differences in the other core parameters. Analysis of variance testing showed statistically significant variation across all core parameters with respect to cell line at 29 DPD (Table S3). Direct pairwise comparisons between cell lines showed that the SCTi003-A line had a significantly lower mean firing rate and mean edge weight relative to H9 and KOLF2.1J, as well as lower mean node degree relative to KOLF2.1J. The H9 line had greater AUNCC relative to H1 and SCTi003-A. The number of single-electrode and network bursts was not significantly different between any pair of cell lines, despite significant variation with respect to cell line across all groups combined. Taken together, these results show that dissociated organoid networks possess inherent variability arising from different organoid batches and donor lines but show a consistent increase in network activity over time with significant temporal differences.

**Figure 2.**
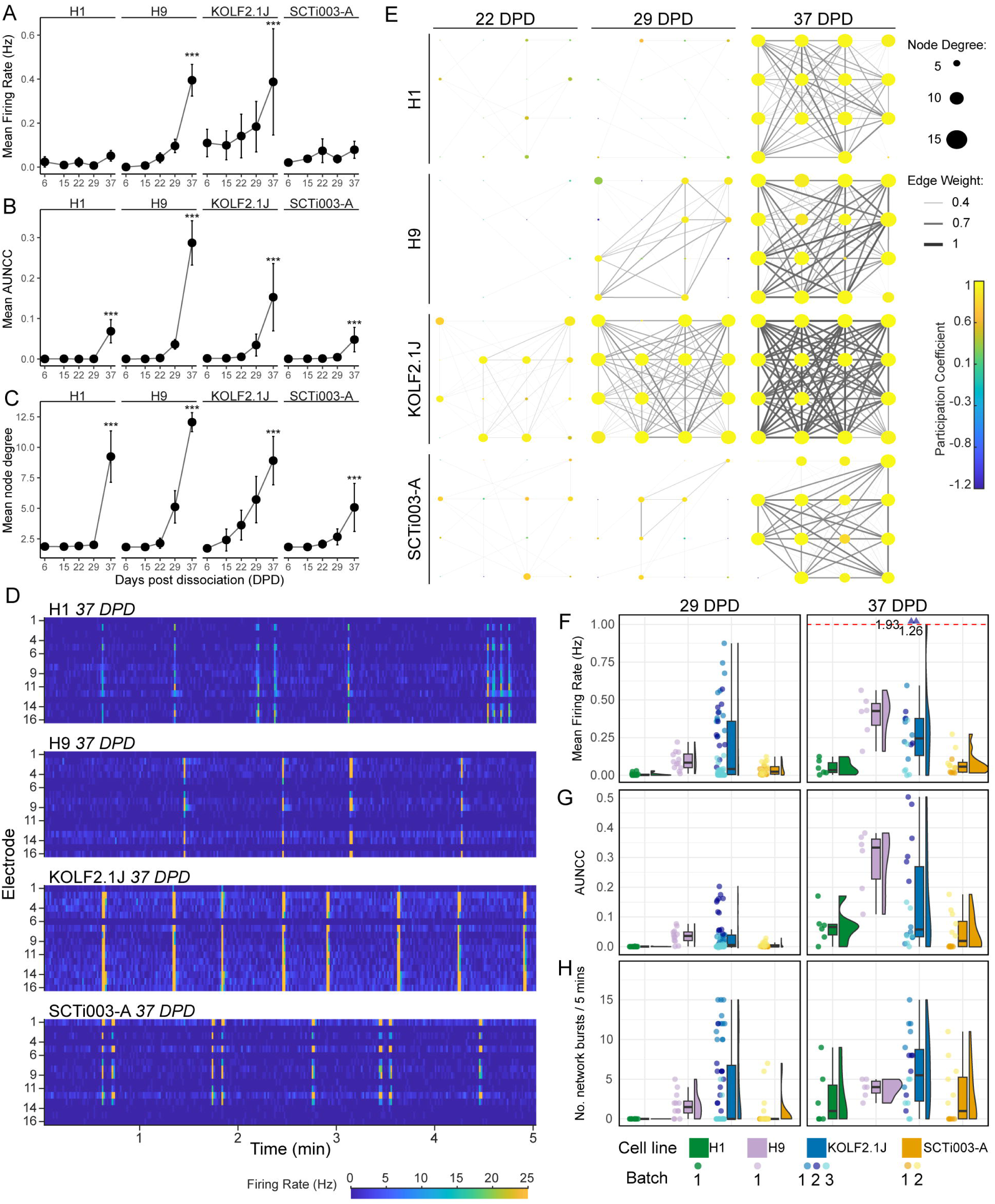
Dissociated organoids from four cell lines develop network connectivity over time. (A-C) Line graphs showing well averages of firing rate (MFR), area under normalized cross-correlation (AUNCC), and node degree at 6-, 15-, 22-, 29-, and 37-DPD. Each point represents the mean of well averages, error bars represent SD. *** indicates p<0.001 for the difference between 6 and 37 DPD. Mixed-effects model with estimated-marginal means post-hoc comparisons for each cell line and DPD combination with Bonferroni correction for multiple comparisons. Full statistical analyses in Table S3. N≥12 wells per line, timepoint, and batch across all lines, with two SCTi003-A batches and three KOLF2.1J batches. At 37 DPD, the N is halved as only control wells are shown, see methods for details. (D) Raster plots at 37 DPD of representative wells with network activity for each cell line showing relative mean firing rate (MFR) in hertz (Hz, color bar) in 1-second bins for each electrode (rows) over 5 minutes scaled to the MFR range in the entire dataset. (E) Scaled network plots showing functional connectivity in at 22, 29, and 37 DPD, corresponding to the wells shown in (D). The nodes (circles) represent neuronal activity at each electrode in the spatial arrangement of the MEA. Node color represents the participation coefficient. Node degree (size of circle) represents the number of functional connections with other nodes. Edges (lines) represent significant functional connections between nodes, and the edge weight (line thickness) represents connectivity strength of connectivity. Participation coefficient color bar, node size, and edge thickness are scaled based on the entire dataset. (F-H) Box (median and interquartile range, IQR) and violin plots of the of the mean firing rate (F), area under normalized cross-correlation (AUNCC) (G), and number of network bursts / 5 minutes (H) per well at two 29 and 37 DPD. Each point represents the well average across 5 minutes of recording. Point colors indicate the batch and cell line. The boxplot color indicates the cell line, ordered from left to right as H1, H9, KOLF2.1J, and SCTi003-A. For clarity, MFR in (F) is capped at 1 Hz (dotted line), two points with MFRs >1 Hz are annotated with their actual values.

### Dimensionality reduction of neuronal and network parameters shows temporal and batch effects in dissociated organoid electrical activity

To integrate electrophysiological and network features across parameters, we visualized the well data for 13 parameters, using a uniform manifold approximation and projection (UMAP) approach across 5 timepoints. The UMAP visualization revealed clustering of wells based on timepoint and batch (Figure 3A- B). Wells with similar activity profiles clustered together, with a progression from low to high firing, burst percentage, bursting, and mean number of significant connections (Figure 3C-F). Separately, we used unsupervised Euclidian clustering to identify clusters based on similarity between parameters (Figure 3G). This clustering showed culture time as the primary factor. Two of the three highest level clusters corresponded to active wells with more mature networks at later timepoints, while the largest cluster consisted of less active wells at earlier timepoints. The proportion of wells in the more active clusters increased with culture time across all lines (Figure 2H). Within these clusters, there were subclusters of wells from the same cell line, suggesting cell line effects. The proportions of wells from each line in each cluster differed by cell line, with KOLF2.1J having greater activity and proportion of wells in the more active cluster at 29 DPD. We further analysed batch and cell line effects at 37 DPD using principal component analysis (PCA) (Figure 3L-O). PCA analysis at 37 DPD suggested a high degree of consistency across batches and cell lines for most experiments (Figure 3M). We saw a batch effect for wells from batch (1) of the SCTi003-A line, which were less active and clustered separately from the remaining wells at 37 DPD, while wells from batch (2) of the SCTi003-A line were more active and clustered with most of the wells from the other cell lines. Reassuringly, two KOLF2.1J line wells that were more active, and which clustered separately from the remaining wells in the PCA, were also independently identified as the separate cluster 3 in our clustering analysis. These results suggest that while batch effects and random differences in networks between wells contribute to variation in dissociated organoid electrophysiological activity, analysis across multiple parameters reveals consistency in activity across batches and cell lines.

**Figure 3.**
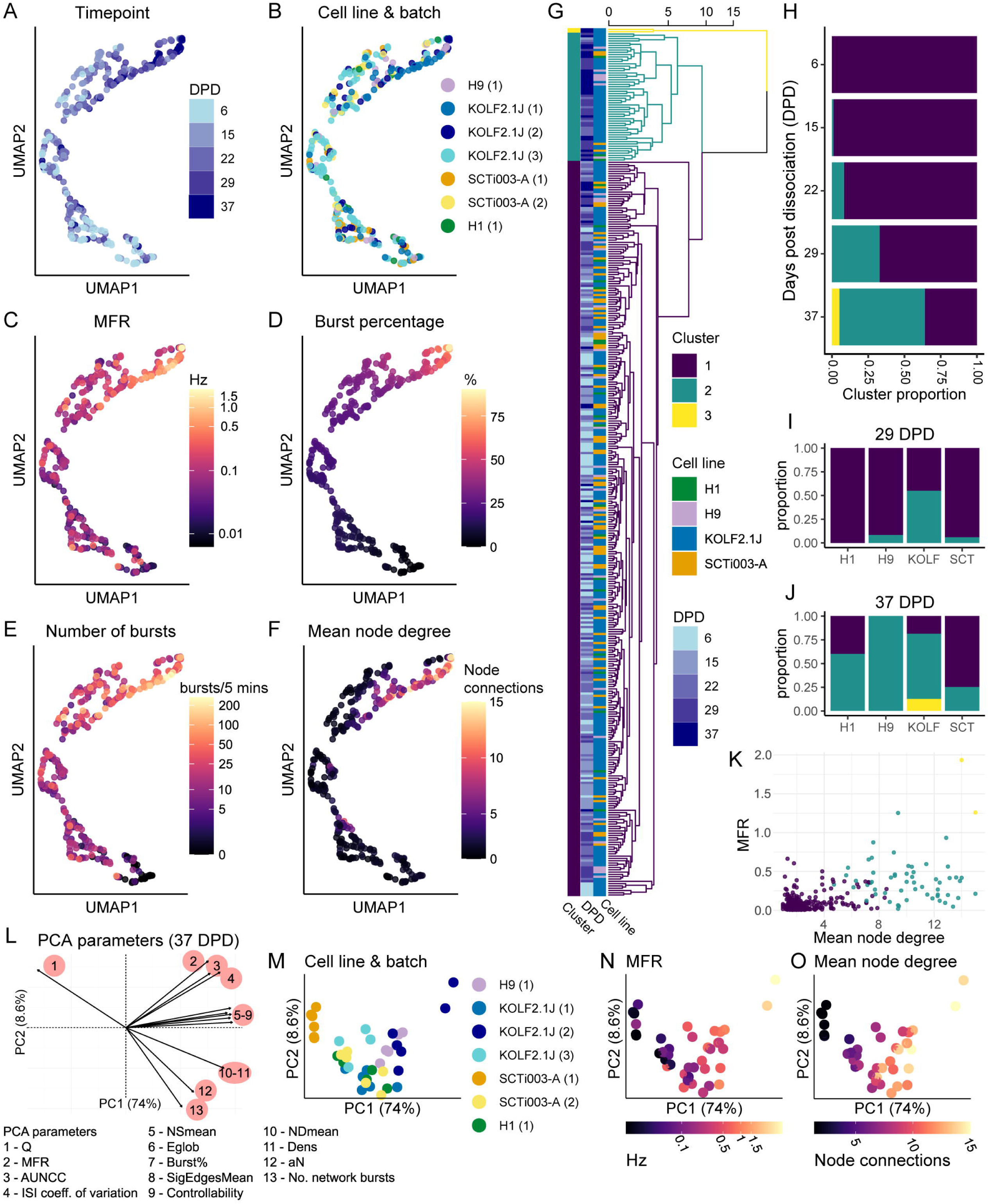
Dimensionality reduction of neuronal and network parameters shows temporal and batch effects in dissociated organoid electrical activity. (A-F) Uniform manifold approximation and projection (UMAP) visualizations of MEA wells from 13 neuronal activity and network parameters showing all wells (points) with active electrodes at 6-, 15-, 22-, 29-, and 37-DPD. Color-coded by timepoint (A), cell line and batch (B), MFR (log10 color scale) (C), burst percentage (D), number of bursts (log10 color scale with 0 shown as black) (E), and mean node degree (F). See also Figure S3. (G) Dendrogram of Euclidean hierarchical cluster analysis using an average linkage method of all wells with active electrodes across five timepoints based on 13 scaled MEA parameters. Branch length indicates degree of similarity in activity between wells. The rightmost link and branches were proportionally compressed to improve visualization. The dendrogram is colored by the top three identified clusters (k=3). The color bars (left) represent the cluster, cell line, and DPD of each well. (H) Bar chart showing proportion of wells assigned to each cluster by timepoint. (I-J) Bar charts of the proportion of wells assigned to each cluster at 29 DPD (I) and 37 DPD (J), grouped by cell line. (K) Scatterplot of mean node degree and MFR for all wells with active electrodes across five timepoints colored by assigned cluster, where each point represents the well average at one timepoint. (L-O) Principal component analysis (PCA) of 13 MEA parameters for all wells with active electrodes at 37 DPD. (L) Biplot showing the impact of each MEA parameter on the top two principal components. The parameters are numbered clockwise and are listed below the plot. PCA plot colored by cell line and batch (M), by MFR (log10 color scale) (N), and by mean node degree (O). Each point in (M-O) represents one well at 37 DPD. Abbreviations: An, network size; AUNCC, area under normalized cross-correlation; Burst%, mean percentage of spikes in bursts; Dens, network density; Eglob, global efficiency; ISI coeff. of variation, inter-spike interval coefficient of variation; MFR, mean firing rate; NDmean, mean node degree; No. network bursts, number of network bursts; NSmean, mean node strength; Q, modularity score; SigEdgesMean, mean significant edge weight.

### Development of spontaneous electrical activity and network properties vary between batches

To further characterize batch effects in dissociated organoids, we analysed the dynamics of acquisition of network activity across three batches of KOLF2.1J dissociated organoids (Figure 4), as this line has been suggested as a reference line across studies^62^. Spontaneous activity was present across all three KOLF2.1J batches, but network bursting only developed over time in two of the batches (Figure 4D).

**Figure 4.**
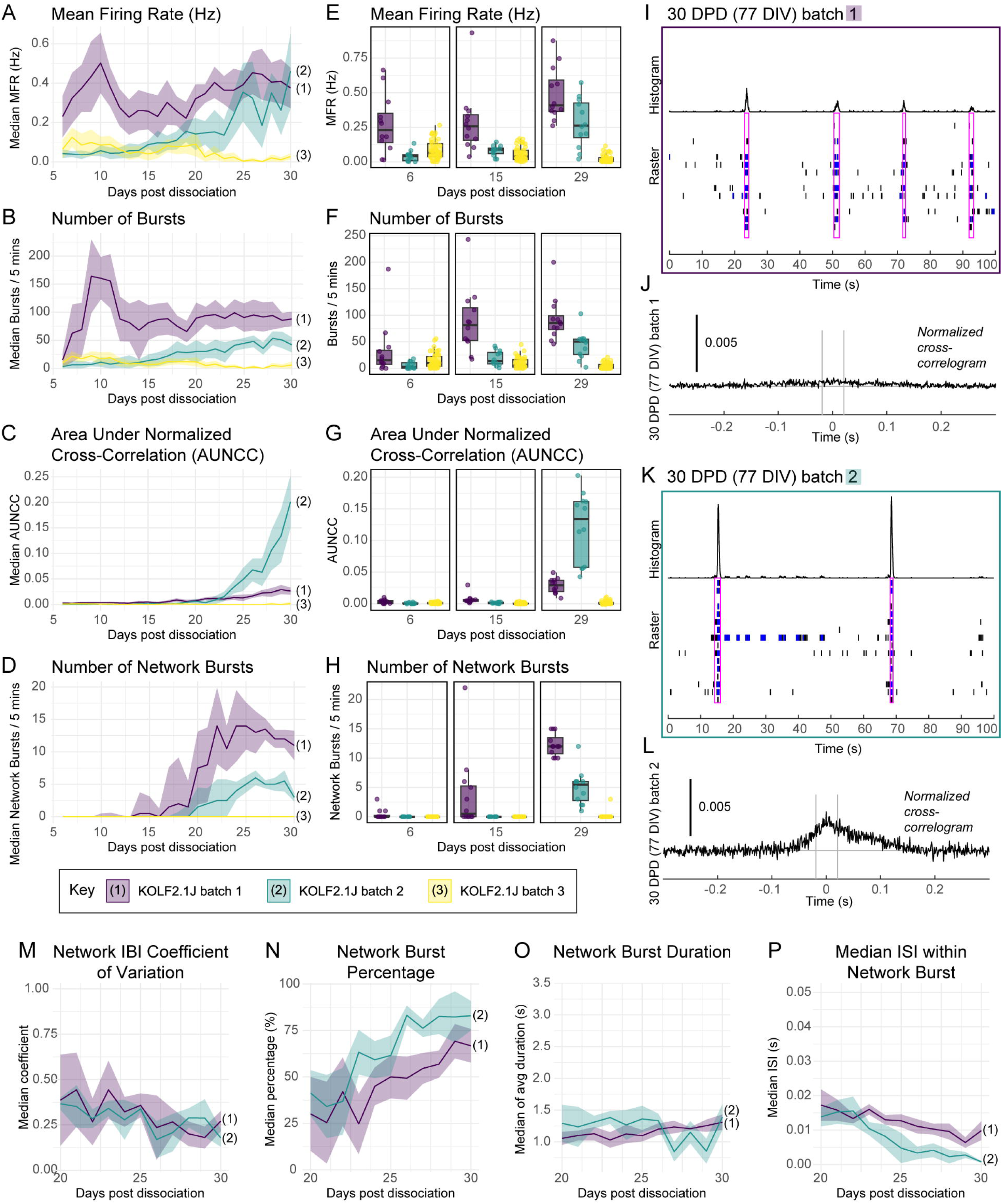
Development of spontaneous electrical activity and network properties vary between batches. (A-D) Line graphs showing the median firing rate (A), number of bursts (B), AUNCC (G), and number of network bursts (H) across three batches of dissociated KOLF2.1J organoids, with one timepoint per day (5 mins recording). The line represents the median of all well averages (N=12 for batch 1 and 2, N=24 for batch 3). The ribbon represents the IQR. Colors indicate batch. (E-H) Boxplots of the min, max, median, and IQR of the MFR (E), number of bursts / 5 minutes (F), AUNCC (G), and number of network bursts / 5 minutes (H) per well at three different timepoints (6, 15, 29 DPD), KOLF21.J line. Points represent well averages across 5 minutes of recording. Colors indicate batch. (I-L) Comparison of two KOLF2.1J dissociated organoid batches with network activity. Raster plots from one representative well of each batch (I, K) showing spikes recorded across 100 seconds at 30 DPD (77 DIV), single electrode bursts (blue), and network bursts (pink boxes). Normalized well-wide cross-correlograms (J, L) for the corresponding wells in (I) and (K) are shown below. The cross-correlogram shows a pooled measure of the relatedness of spike trains across all possible electrode pairs in the well. A higher peak indicates that spikes from electrode pairs occur closer together in time. The time axis indicates the time lag from 0 for each spike in a paired electrode relative to the reference spike in the corresponding electrode, across all electrode pairs. The gray lines indicate the synchrony window. (M-P) Line graphs of parameters for two KOLF2.1J dissociated organoid batches with network activity, showing the median of all well averages for network inter-burst interval (IBI) coefficient of variation (M), network burst percentage (N), network burst duration (O), and median/mean inter-spike interval (ISI) within network bursts (P) between 20-30 DPD. The ribbon shows the IQR. N=12 wells per batch.

Batch differences were present not just in the absolute level of firing and network bursting, but also in the dynamics over time (Figure 4 A-D). While network bursting and synchrony, if at all present, increased with time after ∼15-20 DPD, bursting and spike parameters had a more complex pattern. In one batch, there appeared to be two peaks of high firing, with a high mean firing rate and bursts early after dissociation (∼10 DPD) and a second peak past ∼25 DPD. In the batch where network bursting did not occur, there was also an early peak in firing at ∼10 DPD. In this batch, spontaneous firing decreased over time but started out higher than in another batch that did develop network activity (Figure 4 A, E). The differences in overall activity levels were not simply due to different numbers of active electrodes as weighted mean firing rate was still the lowest in batch 3 and highest in batch 1 (Figure S4).

Interestingly we also observed differences in the characteristics of network burst events. The KOLF2.1J batch with the highest AUNCC after 25 DPD (batch 2, Figure 4C) had less frequent network bursting (Table S2). The differences in synchrony suggested differences in the characteristics in network burst events, which we further characterized at 30 DPD (Table S2). This difference in network burst characteristics was also apparent when comparing raster plots of activity from representative wells (Figure 4I, 4K). Other parameters were more consistent over time between the three batches (Figure S4) and the variation between wells in these parameters (Figure 4) was similar between the two batches that had synchronized network activity (Figure S4). Some parameters of network activity were similar between the two batches with network activity, such as network IBI coefficient of variation, burst percentage, network burst duration, and median/mean ISI within network bursts (Figure 4 M-P).

Overall, dissociated organoids show clear batch differences at the level of individual parameters, with some parameters, such as firing rate, showing greater variability than others.

### Dissociated organoid culture morphology, cell type and regional identity are potential contributors to variation in MEA parameters between batches

Given the variability in spontaneous activity between different cell lines and batches, we predicted that this difference could be due to differences in cell and regional identity composition between experiments. We used immunocytochemistry of cells plated from the initial dissociation grown in parallel to the MEA experiments and bulk RNA-seq of RNA extracted from the MEA plate after the final recording at 37 DPD to characterize the degree of variation in regional identity and cell type composition in the dissociated cultures (Figure 5). We qualitatively compared cells from dissociated organoids from a highly active batch (KOLF2.1J batch 2), medium active batch (SCTi003-A batch 2), and less active batch (H1 batch 1), (Figure 5A-C). We stained cells for markers of excitatory and inhibitory neurons, glia, and dorsal forebrain.

**Figure 5.**
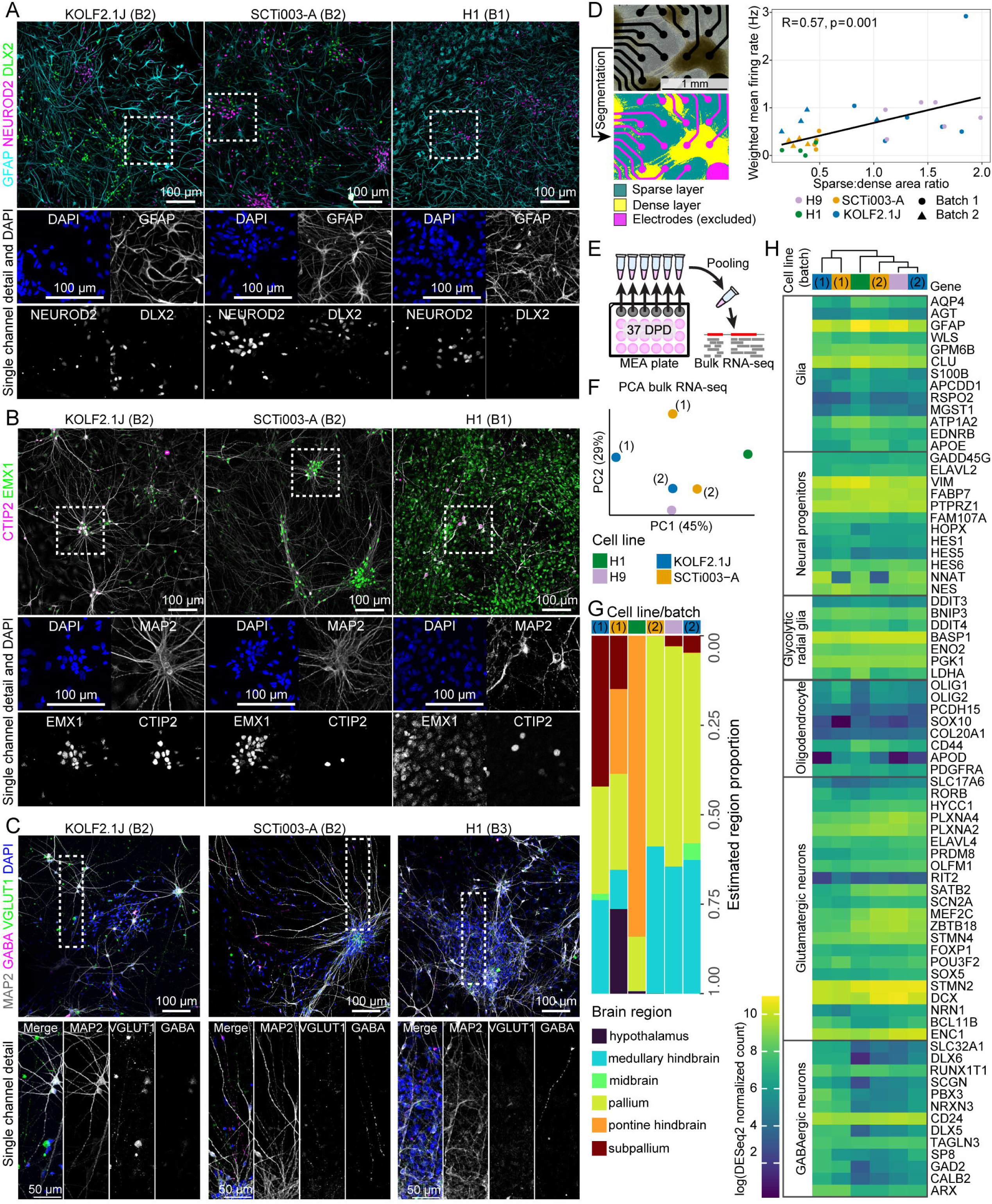
Dissociated organoid culture morphology, cell type and regional identity are potential contributors to variation in MEA parameters between batches. (A-C) Immunocytochemistry (ICC) for excitatory and inhibitory neuronal markers and the glial marker GFAP in cells from dissociated organoids grown on coverslips in parallel to MEA experiments at 37 DPD (84 DIV). Brightness and contrast adjusted for clarity. Cells from a highly active batch (KOLF2.1J batch 2, left), medium active batch (SCTi003-A batch 2, middle), and less active batch (H1 batch 1, right). Regions indicated by the box (white dotted line) are shown enlarged below each main image, with DAPI (blue) and individual channels shown in grayscale. Immunostaining for GFAP, NEUROD2, DLX2 (A), for MAP2, EMX1, CTIP2 (B), and for VGLUT1 and GABA. (D) Analysis of cell clustering in dissociated cultures and correlation to weighted MFR. The colored images show the output of Ilastik-based segmentation into sparse regions (blue), cluster regions (yellow), and electrodes (magenta). A ratio of clustering was calculated as the ratio of sparse area to dense area. The brightfield image (top left) is a well from KOLF2.1J batch 1, at 37 DPD. Dot plot of correlation of cluster ratio at 37 DPD (x axis) compared to weighted mean firing rate (Hz) at 37 DPD. Points represent wells. Point shape indicates batch and color indicates cell line. Statistical analysis conducted using Pearson’s correlation test, N=28 wells. The green line represents the linear regression best fit. (E) Schematic of pooling RNA from all wells within a cell line and batch from an MEA plate for bulk RNA-seq. (F) PCA plot of the 500 most variable genes in the RNA-seq data. The first and second principal components are plotted, where each point represents one sample of pooled wells, point color indicates the cell line, brackets indicate the batch number if applicable. (G) Estimated representation of brain structures in dissociated organoids based on deconvolution of bulk RNA-seq data using Allen Brain Institute in-situ hybridization (ISH) reference data. Each column represents one sample, with colors representing the proportion of each brain region. Column annotations indicate the sample cell line (color), and batch (brackets). (H) Heatmap of expression of genes (row annotations, right) from cell type signatures (row annotations, left). Genes and categories adapted from Hendriks et al. ^92^. The color scale represents the log(DESeq2 normalized counts), with yellow representing the highest expression. Column annotations indicate the sample cell line (color), and batch (brackets). The dendrogram (top) represents the hierarchical clustering of the samples.

The dissociated organoids consisted of a mixture of cell types, with star-shaped GFAP-positive cells, presumably astrocytes, MAP2-positive cells with a neuronal morphology, of which some nuclei were positive for the excitatory markers CTIP2 and NEUROD2, or inhibitory marker DLX2 (Figure 5A-B). These neurons were found in mixed clusters together. In addition, there were processes with distributed puncta of VGLUT1 and separate, non-overlapping GABA-positive processes (Figure 5C), which appeared to radiate out of the same neuronal clusters. Although all three cultures contained cells that were positive for the markers used, there were some clear differences in composition in the dissociated culture from H1 organoids, with what morphologically appeared to be an increased number of progenitors, increased number of GFAP-positive cells, and more diffuse and less intense EMX1 expression (Figure 5A-B). The H1 culture contained fewer MAP2-positive neurons and appeared denser in terms of overall number of DAPI-positive nuclei.

Given the variability in activity between wells in the same experiment, we hypothesized that well-to-well differences in spontaneous activity may also be due to the random differences in the sizes and positioning of clumps in different wells that developed over time. We quantified the degree of clustering in different wells as a cluster ratio of sparse to dense areas and plotted this against firing rate at active electrodes (Figure 5D). There was a significant positive correlation, where the sparser the well, the greater the mean firing rate was (R=0.57, p=0.001). However, this appeared to be cell line or batch dependent. Batches with high cell density had very low firing rates, but batches with clustering and sparse areas had more variable firing rates. We did not see significant correlation of cell clustering with bursting, network bursting, or AUNCC, suggesting that network characteristics are less dependent on cell density than firing rate. There were still clear contributions of cell line and batch (Figure 5D).

Deconvolution of bulk RNA-seq data^63^ showed that all dissociated organoid samples were estimated to contain cells representing pallium, however, the proportion of pallium varied. Some samples showed reduced pallial representation with other brain regions, including pontine hindbrain, subpallium, midbrain, and hypothalamus, present at varying proportions (Figure 5G). Notably, the H1 line dissociated organoids, which had the lowest firing rates, were estimated to primarily consist of prepontine hindbrain cells. Batch 1 of the SCTi003-A line, which clustered separately at 37 DPD based on MEA activity parameters (Figure 3M) also clustered separately based on gene expression (Figure 5F) and had increased proportion of hypothalamus (Figure 3G). To further characterize the cell type composition of the dissociated organoid samples, we examined cell type marker expression (Figure 5H). Consistent with the ICC data, we found that the H1 line culture had altered levels of cell type markers, such as increased GFAP expression. Hierarchical clustering based on the selected cell type markers mirrored PCA clustering based on the top 500 most differentially expressed genes between the samples, with batch 1 of KOLF2.1J and SCTi003-A clustering separately (Figure 5H).

This analysis shows that differences in electrical activity parameters are linked to consistent differences in cell identity and cell type composition, which can be detected at both the protein and gene expression levels.

### Dissociated organoids respond to a chemical plasticity stimulus

To test whether the neurons in the dissociated cultures showed plasticity in response to a stimulus, we used a chemically induced LTP-like (chLTP) stimulation paradigm^50^ (Figure 6A). Following chLTP, there was an initial decrease in spontaneous firing 30 minutes after treatment. Three hours after treatment, mean firing rate, number of bursts, AUNCC, and number of network bursts increased relative to baseline in chLTP-treated wells (Figure 6). The greatest response to chLTP occurred between 6-24 hours and then diminished. At 6 hours after treatment, there was a significant increase in mean firing rate, AUNCC, number of bursts and network bursts in the chLTP condition relative to vehicle (Table S3). There was an increase in single electrode bursting and synchrony, with shorter bursts compared to baseline measurements or the vehicle condition (Figure 6 B, C). While there was not a significant increase in node degree and edge weight, we observed increased strength of network connections in more active wells (Figure 6D). PCA analysis across the six core parameters showed a clear shift in the overall activity profile of chLTP-exposed wells; at baseline, wells from both conditions clustered together (Figure 6E), while at 6 hours post chLTP, there was clear separation of wells based on whether they were treated with chLTP or vehicle (Figure 6F). The magnitude of response to chLTP differed between replicates and wells (Figure 6G-L). Across both replicates, median single-electrode measures of activity, firing rate and number of bursts, increased, although the increase was larger in the first replicate. In contrast, synchrony and number of network bursts increased minimally in the second replicate (Figure 6 I-J). Some wells did not have a large response to chLTP, however in wells with bursting, there was a consistent decrease after chLTP in the variability of the ISI, meaning increased consistency of firing, and a reduction in the inter-burst interval, meaning more regular bursting. These two parameters clearly separated the vehicle and chLTP conditions (Figure 6 K-L). Thus, despite variability in responses between wells and replicates, dissociated organoids show plasticity in response to stimulation, with chLTP being a reproducible method for increasing network activity in dissociated organoids.

**Figure 6.**
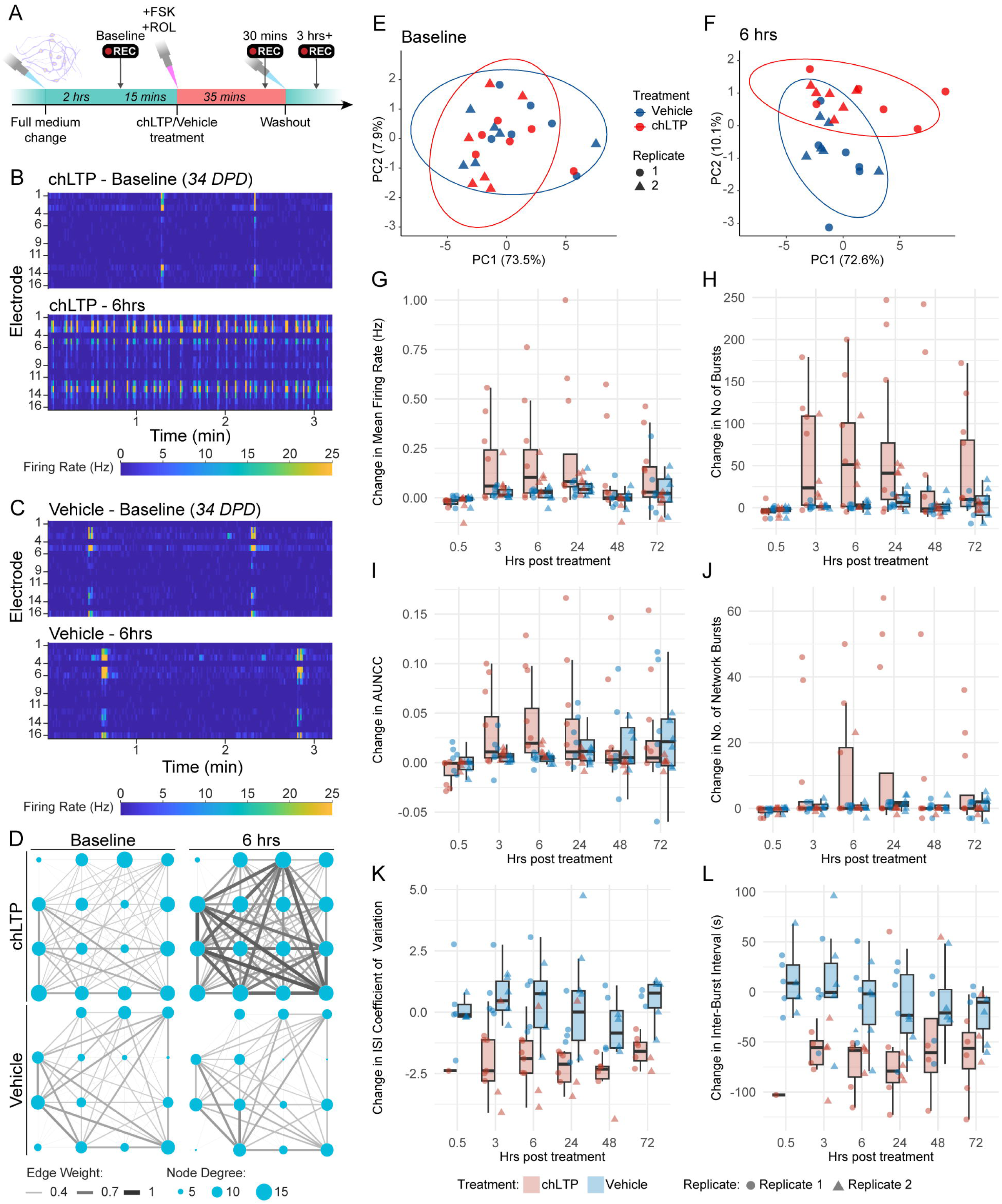
Dissociated organoids respond to a chemical plasticity stimulus. (A) Schematic of chLTP treatment and data collection timeline. (B-C) Raster plots of firing rate at 34 DPD for two representative wells at baseline and 6 hours following chLTP (C) or vehicle (D). The raster plots correspond to the two wells and timepoints shown in (B). The raster plots show MFR in hertz (Hz, color bar) in 1-second bins for each electrode (rows) over 3 minutes scaled to the range of MFR in the entire dataset. (D) Scaled network plots showing functional connectivity in representative wells rate at 34 DPD for two representative wells at baseline (left) and 6 hours following chLTP (top right) or vehicle exposure (bottom right). The nodes (circles) represent the neuronal activity observed from neuron(s) at each electrode in the spatial arrangement of the MEA. The node degree (size of circle) represents the number of functional connections with other nodes. The edges (lines) represent significant functional connections between nodes, and the edge weight (line thickness) represents the strength of connectivity. (E-F) PCA analysis of 13 MEA parameters measured at baseline (E) and 6 hours following chLTP (F). PCA plots of the first and second principal components. Each point represents one MEA well, colored by treatment. Point shape indicates the replicate. The data ellipses represent the 95% confidence interval, expected to enclose 95% of bivariate-normal t distributed data for each treatment condition. (G-L) Boxplots of difference relative to baseline in MEA parameters after chLTP or vehicle treatment. Boxplots show the min, max, median, and IQR of the change in MFR (G), number of bursts / 5 minutes (H), AUNCC (I), number of network bursts / 5 minutes (J), ISI coefficient of variation (K), and inter-burst interval (L) per well, at six different timepoints (0.5, 3, 6, 24, 48, 72 hours post treatment). Points represent the mean of one well across 5 minutes of recording. Point color indicates treatment (vehicle blue, chLTP red), shape indicates the replicate. In (K-L), only wells with bursts present are shown. Measured in KOLF2.1J line dissociated organoids, with treatment starting at 34 DPD (replicate one) and 29 DPD (replicate 2).

### 17β-estradiol does not affect electrical activity in dissociated organoids

Given the known role of 17β-estradiol in enhancing LTP in rodents^51–57^, we conducted a pilot experiment, where we tested the effect of E2 in combination with chLTP. Overall, we did not detect a significant effect of E2 (Figure S5, Table S3). E2 by itself did not affect any of the parameters in both replicates and was not distinguishable from the vehicle condition. In line with this, there did not appear to be any separation of the E2 pretreatment and vehicle wells, nor separation of the chLTP and chLTP+E2 wells across both replicates in PCA analysis (Figure S5E). Hierarchical clustering on the PCA data showed that all but one of the samples without chLTP clustered together (Figure S5F). The chLTP treated samples clustered into three groups, presumably based on the magnitude of response to chLTP.

To determine whether longer-term E2 exposure affects the development of spontaneous electrical activity in dissociated organoids, we exposed dissociated organoids to 10nM E2 or vehicle daily for seven days (Figure S6A). Overall, there was no consistent effect of E2 exposure across both batches of the KOLF2.1J line when compared to vehicle at any timepoint (Figure S6). Although there continued to be significant batch differences and changes over time in parameters such as number of network bursts, there was no difference between E2-exposed and vehicle wells. For example, the number of network bursts was higher on day 8 relative to baseline in batch 2 (Figure S6 E, I), reflected in a significant effect of day and batch (Table S2), but this increase was the same across vehicle and E2-exposed wells.

Similarly, we did not see any effect of treatment for the H9 and SCTi003-A cell lines. PCA analysis of wells across all three cell lines and the two KOLF2.1J batches showed a clear batch/cell line effect, but did not show any obvious treatment effect at either timepoint (Figure S7). Together, our analysis of dissociated organoids acutely exposed to chLTP, chLTP with E2, E2 alone, or exposed to chronic E2, showed that while chLTP induced changes in spontaneous activity across multiple parameters, E2 did not appear to have an effect under the conditions tested.

## DISCUSSION

Neuronal networks have been characterized in limited numbers of organoids and donor lines ^43–45^. Building on existing studies, we characterised the development of neural networks specifically in unguided dissociated organoids, using multiple cell lines and batches. We observed a gradual development of activity over time in the dissociated organoids, in line with other studies done in 2D and 3D neuronal networks ^23,44,45,64^. More complex activity generally arises with longer culture periods– studies using organoid slices have shown long distance connectivity at 130 days of culture ^46^, while whole unguided organoids had network bursting at 120 days, with weak activity at one month of culture ^45^. Dissociated organoids developed spontaneous network activity much earlier in several replicates, with network bursting in some replicates occurring after around 20 DPD. This is similar in timing to the synchronised calcium bursts at 30 DPD observed by Sakaguchi *et al.* ^43^ in dissociated organoid cultures.

In a study using iPSC neurons from multiple lines, Mossink *et al.* ^20^ only saw subtle variability between wells and experiments and no line-specific differences, recommending a minimum of two replicates per line. However, this would be expected with the much more homogeneous monoculture model used. We observed a hierarchy of variability at the level of cell line, batch, and variability between neural networks in each well. Like Mossink *et al.* ^20^, we observed variability in some parameters but not others, including similar network burst duration and similar magnitudes of within-batch variability for parameters such as inter-burst interval. We did not see any sex differences or systematic differences between iPSC and ESC lines, in agreement with Mossink *et al.* ^20^. We suggest that care needs to be taken in selection of MEA parameters for analysis, in particular, in line with others^20^, we saw that mean firing rate is highly variable and is dependent on neuronal density. Although neuronal clustering affected firing rate, the same phenomenon has been observed in other dissociated organoid cultures^43^ and may accelerate the development of network activity.^65^ Our integration of multiple parameters using a dimensionality reduction approach, similar to what has been done with neuronal monocultures^20^, may be a useful method for extracting electrophysiological phenotypes while avoiding the noise associated with looking at single parameters like mean firing rate.

In our assessment of the cell type composition in dissociated organoid cultures, we showed that dissociated organoids contain a mixture of inhibitory and excitatory neurons. iPSC GABAergic neurons can synaptically integrate with glutamatergic neurons in co-cultures, resulting in reduced firing, network bursting, and network burst duration compared to cultures without GABAergic neurons ^66^. Increasing the proportion of GABAergic neurons relative to glutamatergic neurons significantly decreased the number of network bursts and firing rate ^66^. Thus, variability in spontaneous activity in dissociated organoids between experiments could be explained by variable proportions of excitatory and inhibitory neurons. Consistent with this, previous whole and dissociated organoid studies have reported that the ratio of glutamatergic GABAergic neurons can affect network activity parameters ^43,45^. Similarly, astrocytes are critical for neuronal plasticity, synapse function and synapse formation^67^. Lower numbers of astrocytes in neuron and astrocyte co-cultures on an MEA have been shown to enhance development of network activity ^65^. In line with this, we saw an apparent difference in the number of GFAP-positive cells in the H1 dissociated culture with minimal spontaneous activity. We suggest that cell type composition should be assessed in organoid MEA experiments containing mixed proportions of excitatory and inhibitory neurons going forward. Future studies could investigate whether dissociated guided organoids are more consistent, relative to unguided, in composition and activity between batches and cell lines.

Previous studies have demonstrated changes in organoid network activity in response to electrical stimulation and drug treatments ^43,64^, although most only looked at acute changes in network activity of less than one hour. Long term potentiation in organoid slices in response to high frequency electrical stimulation on an MEA was reported by Zafeiriou *et al.* ^64^ in the form of a consistent increase in firing following stimulation. In dopaminergic and cortical neurons co-cultured with astrocytes, Pré *et al.* ^50^ observed increased firing rate and network burst frequency following chLTP that peaked between 4-24 hours after treatment and lasted up to 72 hours. We observed similar timing in the chLTP response, with increased firing and network bursting, which peaked between 6-24 hours. Pré *et al.* ^50^ also showed variation in the magnitude of response between eight independent experiments. We observed similar variability between experiments in the dissociated organoids, with a consistent response across parameters such as ISI coefficient of variation and inter-burst interval. The coefficients of variation for these parameters were also less variable at baseline. Thus, with careful parameter selection^20^, chLTP can be extended for use in more complex cultures and cerebral organoids. We found no effect of E2 on spontaneous activity. Rodent studies have suggested that E2 induces silent synapses, which increase the response to subsequent stimuli ^68,69^. One recent study using human iPSC neurons found that E2 at a slightly higher, less physiological^70^ concentration (100nM) induced changes in both the amplitude and spiking frequency of calcium oscillations, although this effect varied between cell lines ^71^. In previous studies in rodents, increased spine density has been associated with functional changes that were detected using patch-clamp intracellular recordings^72^ where in some cases action potential amplitude was unaffected^68^. Use of electrical stimulation and patch clamping could be used in dissociated organoids to test the effects of E2.

In summary, we showed here that cerebral organoids dissociated to 2D cultures consistently develop neural networks over time. While sex or line type (iPSC or ESC) did not appear to affect the development of network activity, there were batch differences in parameters that we attribute to differences in cell composition and ratio in the cultures. We suggest that in future studies, this variation could be exploited by characterising neural networks, cell type composition, and gene expression in each well to enhance our understanding of neural network development. Finally, we showed using a chemical stimulation paradigm that the dissociated organoid networks are capable of a long-term plastic response to a stimulus. These results show that cultures of dissociated cerebral organoids can be used similarly to primary neuronal cultures, with important implications for disease modelling and pharmacological screening, offering a platform to study patient-derived neuronal networks and test the effects of genetic or environmental perturbations on network connectivity.

## ACKNOWLEDGMENTS

The authors acknowledge funding support from UK Medical Research Council, Grant Nos. MR/L021064/1 (D.P.S.), MR/Y012968/1 (A.C.V., D.P.S.), MR/X004112/1 (D.P.S.), MR/Y008170/1 (D.P.S.), and MR/Y012968/1 (A.C.V., D.P.S.) and from The Simons Foundation Autism Research Initiative (SFARI). D.P.S. is also a recipient of an Independent Researcher Award from the Brain and Behavior Foundation (formally National Alliance for Research on Schizophrenia and Depression) (Grant No. 25957). A.P. is in receipt of the MRC- Institute for Translational Neurodevelopment (ITND) Ph.D. studentship, as part of the MRC Centre for Neurodevelopmental Disorders (Medical Research Council MR/P502108/1). A.P. and D.P.S. also acknowledge funding from Psychiatry Research Trust. The authors acknowledge use of King’s Computational Research, Engineering and Technology Environment (CREATE). The authors thank George Chenell of the Wohl Cellular Imaging Centre (King’s College London) for technical support, Niamh O’Brien for advice on organoid dissociation methods, and Iva Kelava for training in cerebral organoid culture.

## AUTHOR CONTRIBUTIONS

Conceptualization, A.P., A.C.V. and D.P.S.; methodology, A.P., A.C.V. and D.P.S.; Investigation, A.P., L.D.P., R.N.; writing—original draft, A.P.; visualization, A.P, writing—review & editing, M.A.L., A.C.V. and D.P.S.; funding acquisition, M.A.L., A.C.V. and D.P.S.; resources, M.L.; supervision, M.A.L., A.C.V., and D.P.S.

## DECLARATION OF INTERESTS

The authors declare no competing interests.

## SUPPLEMENTAL INFORMATION

Document S1. Figures S1–S7 and Tables S1-S2 Table S3. Full statistical test results

See table S1 for parameter definitions.

## METHODS

### EXPERIMENTAL MODEL AND STUDY PARTICIPANT DETAILS

#### Cell line information

Two human induced pluripotent stem cell (hiPSC) lines, SCTi003-A (XX) and KOLF2.1J (XY) and two human embryonic stem cell (hESC) lines, H9 (XX) and H1 (XY) were used for experiments. To ensure high quality lines with respect to low passage number, characterisation, and standardisation, cells for the project were sourced directly from stem cell banks under material transfer agreements (MTA). The SCTi003-A line was obtained from Stem Cell Technologies (200-0511, Lot# 2206422017) as passage 32. The KOLF2.1J line was obtained from Jackson Laboratory as passage 4. The H9 (WA09) and H1 (WA01) lines were obtained from WiCell as passage 29 (Lot# WB67619) and passage 30 (WB34445), respectively. All cell lines were derived from control patients with no disease diagnosis. All cell lines had received full consent and ethical approval for research use. Approval for use of hESC lines in this study was obtained from the UK Stem Cell Bank (UKSCB) Steering Committee (reference number SCSC22- 03).

| CELL LINE | HPSCREG ID | TYPE | SOURCE TISSUE | REPROGRAMMING VECTOR | SEX | AGE | ETHNICITY |
| --- | --- | --- | --- | --- | --- | --- | --- |
| H9 | WAe009-A | hESC | Blastula (ICM) supernumerary embryos from IVF treatment | N/A | Female | N/A | White |
| H1 | WAe001-A | hESC | Blastula (ICM) supernumerary embryos from IVF treatment | N/A | Male | N/A | White |
| SCTi003-A | SCTi003-A | iPSC | Blood | Non-integrating (proprietary) | Female | 45-49 | White |
| KOLF2.1J | WTSli018-B-12 | iPSC | Fibroblasts | Non-integrating (Sendai virus) | Male | 55-59 | White |

All cell lines were expanded and banked, and working stocks were generated from banked vials. Medium from all cell lines was submitted for regular PCR-based mycoplasma testing. Stem cells were stored in liquid nitrogen suspended in freezing medium (10% DMSO in Stemflex medium (Gibco; A3349401)). All stem cells were grown on 6-well plates (NUNC) coated with Geltrex (Geltrex LDEV-Free Reduced Growth Factor Basement Membrane Matrix, Gibco A1413201) diluted 1:100 in DMEM (Sigma; D6421). The stem cells were thawed in a 37°C water bath and the freezing medium diluted with DMEM. The cells were then pelleted at 900 RPM for two minutes and resuspended in Stemflex medium with 1:100 Revitacell (Gibco; A2644501). The cells were kept in hypoxic conditions at 37 °C and 5% CO2, 5% O2. After 24 hours, the medium was replaced with Stemflex medium, and full media exchanges were performed every 48 hours. The stem cells were passaged once 70-80% confluent by washing with HBSS and then incubating for 3 minutes in room temperature Versene (Gibco; 15040066). The Versene was aspirated and replaced with Stemflex medium, the cells were dislodged by pipetting medium over the cells. The cells were then added to the appropriate volume of medium for splitting into new Geltrex-coated plates (between 1:2 to 1:6). The stem cells were monitored daily using brightfield microscopy (EVOS XL). Wells with 70-80% confluence and no visible signs of spontaneous differentiation were used to generate cerebral organoids. iPSCs used to generate organoids were not allowed to exceed passage 35, and ESCs were not allowed to exceed passage 48.

In addition to cell line quality checks conducted as part of cell line sourcing additional checks for pluripotency and common karyotypic abnormalities were done following established guidelines ^73^. A subset of undifferentiated stem cells from the working stock was grown on glass coverslips and ICC was conducted to check expression of the pluripotency markers NANOG, POU5F1, and PODXL. DNA was extracted from organoids from the highest passage batches of all the cell lines using a Genomic DNA Extraction Kit (Monarch; T3010). The DNA samples were tested for the most common genetic abnormalities that can arise during stem cell passaging and expansion using the hPSC Genetic Analysis Kit (Stem Cell Technologies; 07550) according to the manufacturer’s protocol.

### METHOD DETAILS

#### Generation of unguided cerebral organoids

Cerebral organoids were generated using the commercially available STEMdiff Cerebral Organoid Kit (Stem Cell Technologies; 08570) and associated protocol (Stem Cell Technologies Document #DX21849). Organoids generated on the same day from the same passage number of stem cells were considered part of the same batch. The organoid kit protocol was modified to allow for long-term culture of cerebral organoids using a published protocol ^74^. The following changes to the manufacturer’s protocol were made: the embryoid body (EB) seeding medium was supplemented with a higher concentration of ROCK inhibitor (50 µM) (as recommended in Giandomenico *et al.* ^74^). Smaller EBs were generated using fewer cells (∼1000 cells per EB) as we found that this worked well for the cell lines used. Geltrex was used instead of Matrigel in all steps of the protocol. In brief, the protocol was as follows: First, embryoid bodies (EBs) were generated by seeding stem cells into a 96-well round-bottom ultra-low attachment plate (Corning; 7007) at 0 days *in vitro* (DIV). The EBs were kept in hypoxic conditions at 37 °C and 5% CO2, 5% O2 until 5 DIV. At 5 DIV, the EBs were transferred to induction medium in a 24-well ultra-low attachment plate (Corning; 3473) and hereafter incubated in normoxic conditions at 37 °C and 5% CO2. At 7 DIV, EBs were embedded in Geltrex and transferred to expansion medium until 10 DIV. From 10 DIV onwards, organoids were maintained in maturation medium consisting of STEMdiff Cerebral Organoid Basal Medium 2 and STEMdiff Cerebral Organoid Supplement E (Stem Cell Technologies; 08571). The organoids were placed on an orbital shaker (Infors; Celltron) in the incubator at 37 °C and 5% CO2, with the shaker set to 57 revolutions per minute. The Geltrex was removed from the encapsulated organoids at ∼12 DIV using a 10 ml stripette. Organoids were grown in 50 mm dishes and fed by a full medium change of 5 ml maturation medium twice a week. The organoids were split (e.g., organoids were divided between two dishes) when the medium turned yellow between feedings. The maturation medium was supplemented with 2:100 Geltrex during feeding from 30 DIV until 50 DIV to promote growth of the cortical plate. Antibiotics were not used in any stage of the protocol. The organoids were grown until 45- 47 DIV when they were dissociated. Quality control checks were done throughout the organoid generation and maturation protocol by regular monitoring of the development of the organoids using brightfield microscopy. Quality criteria were based on the recommendations provided by Stem Cell Technologies and additionally in the published protocol ^46^ – these were as follows: Stem cells without any obvious signs of differentiation were used and if EBs failed to form by day 3 of the protocol, the experiment was terminated, and a new vial of stem cells was used. Additional quality control steps included checking that the EB borders showed clearing prior to induction, checking for the formation of radially organized neuroepithelium following induction, checking for formation of neurepithelial buds during expansion, and removal of any organoids showing excessive cyst formation, migratory cells, or disintegration.

#### Dissociation of cerebral organoids for culture on MEA

Whole organoids were dissociated at 45-47 DIV using the Neural Tissue Dissociation Kit (P, Papain) (Miltenyi Biotec; 130-092-628). For each batch and cell line, six organoids from the same batch were transferred to a petri dish (Thermo; 150462) containing 5 ml HBSS with calcium and magnesium (+/+, Gibco 14025-092) using a cut 1 ml pipette. The organoids were then cut into several pieces using a sterile scalpel (Swann-Morton 0501), visually targeting the cortical lobe regions of the organoids. The cut organoids were gently washed in the petri dish using circular movements three times and were then transferred to a 15 ml falcon tube and the supernatant was aspirated. A volume of 2 ml of preheated enzyme mix 1 (50 µl Enzyme P into 1900 µl Buffer X) was added to the organoids to initiate papain-based digestion. The organoids were incubated at 37 °C for 15 mins. Enzyme Mix 2 (DNAse mix containing 20ul of Buffer Y and 10ul of Enzyme A) was then added to the reaction. The organoids were triturated approximately 20 times using a cut 1000 µl pipette tip and then a standard 1000 µl tip was used to break up the organoid pieces. This process was repeated every 5 mins of incubation up to a total of ∼3 times until the solution was cloudy and no tissue pieces or aggregates were observable. Any remaining clumps were removed by filtering through a 40 µm nylon mesh sterile strainer (Fisher brand; 22363547). The strainer was pre-wetted with HBSS without calcium and magnesium (-/-, 1 ml, Gibco; 14170-070) prior to the addition of the organoid cell suspension. The filter was then washed with 2 ml of HBSS (-/-). The cells were pelleted in a centrifuge, the supernatant aspirated, and the cells resuspended in 5 ml of HBSS (-/-) and pelleted again. The cells were then resuspended in FBS recovery medium with 100 µg/ml laminin to achieve a concentration of 7.5 million cells/ml. FBS recovery medium consisted of 445 ml high-glucose DMEM supplemented with GlutaMAX (Gibco; 10569010), 50 ml FBS (Gibco; 10270-106) and 5 ml of 50% (wt/vol) glucose solution ^74^ without antibiotic-antimycotic.

#### Dissociated organoid plating and MEA culture

MEA plates (Axion; M384-tMEA-24w) for culturing cells from dissociated organoids were coated with 0.1% Poly(ethyleneimine) solution (PEI) for 60 minutes at 37°C, 5% CO2. The PEI was deposited on the area of the electrodes as 10 µl droplets. The PEI was rinsed with 200 μl of sterile DI water (ddH2O) 4 times, and the plate was allowed to air-dry overnight. The MEA plate was then coated with 9.6 ug/mL laminin (Sigma; L2020-1MG) in DMEM (Sigma; D6421) for 4 hours at 37°C, 5% CO2 and then rinsed three times with HBSS (-/-). A volume of 10 µl of cell suspension (approximately 75,000 cells per well) was applied to the recording electrodes on the electrode array and the plated cells were incubated 60 minutes at 37°C, 5% CO2. FBS recovery medium was then slowly applied to the wells to a final volume of 0.5 ml. Half media changes were performed every other day with BrainPhys+SM1 (BrainPhys (StemCell Technologies; 05790) supplemented with 200 µl/9.8 ml NeuroCult SM1 (StemCell Technologies; 05711)). The BrainPhys+SM1 medium was also supplemented with 2 ug/ml laminin weekly.

#### MEA recording

Continuous MEA recordings were made across all electrodes in real time using the Axion Maestro and Axis Navigator software (12.5 kHz sampling rate, 200-3000 Hz band-pass) and were stored as .raw files. The temperature and CO2 were allowed to stabilise at 37°C and 5% prior to recording after inserting the plate into the Maestro. Recordings were always made prior to any media changes, or after a minimum of two hours after a medium change if required by the treatment protocol. For acute treatment experiments, a 5-minute baseline recording was taken prior to adding the treatment. Recordings were made over 5- minutes and subsequently analysed. For comparison of spontaneous activity between cell lines and batches of dissociated organoids from each cell line, daily five-minute recordings were taken from 6 DPD to 37 DPD in one batch of H1 dissociated organoids, one batch of H9 dissociated organoids, two batches of SCTi003-A dissociated organoids, and 3 batches of KOLF2.1J dissociated organoids. Recordings consisted of N=12 wells per batch and cell line, except for the third batch of KOLF2.1J organoids, where 48 wells were used, split across two plates. Half of the wells were exposed to 17β-Estradiol from 29 DPD onwards, control recordings from 30 DPD onwards thus consisted of N=6 wells per batch and cell line.

#### Treatment experiments

##### 17β-Estradiol Treatment

17β-Estradiol powder (E2758-250MG; Sigma) was dissolved in 1 ml absolute ethanol and a 20 μg/ml stock solution made by adding DMEM/F-12 phenol red-free (PR-F) (Gibco; 21041025). Vehicle stock solution was made in the same manner, but without 17β-Estradiol powder. The 17β-Estradiol and vehicle stocks were stored as single-use aliquots at -20 °C for up to 1 month. Diluted treatment solutions were prepared freshly each day, by sequential dilution of the stock, first to a 10 µM solution in DMEM PR-F, then to a working stock of 1 µM in BrainPhys+SM1 PR-F. 1 µM 17β-Estradiol or vehicle solution was added to PR-F cell culture medium, with a final concentration of 10 nM. Dissociated organoids were transitioned to medium without phenol red at least 2 days prior to commencing treatment experiments.

17β-estradiol or vehicle were spiked directly into the dissociated cultures every day following recording (6 µl spike to 600 µl of medium in each 24-well plate).

##### chLTP

chLTP treatment was conducted according to the method described by Pré *et al.* ^50^ for iPSC neurons. The chLTP experiment was done using KOLF2.1J dissociated organoids from one batch of nine pooled organoids in two replicates on two different days, with 6 wells per condition per replicate (12 total wells for chLTP and 12 vehicle wells). The chLTP treatment consisted of Rolipram (Tocris; 0905/10), stock prepared as 25 mM in DMSO, and Forskolin (Tocris; 1099/10), stock prepared as 50 mM in DMSO. On the day of treatment, a full medium change with 600 µl media per well was done two hours prior to the start of the treatment. Next, a baseline recording was taken and 17β-estradiol to a 10 nM concentration or vehicle (containing the same concentration of ethanol) were added for 15 minutes. After 15 minutes of pre-treatment, 30 µl medium was removed from each well of the MEA and 30 µl of 20x chLTP treatment solution was added, consisting of 1000 µM Forskolin and 2 µM Rolipram diluted in Brainphys media. The vehicle condition treatment consisted of the same amount of DMSO in Brainphys, but without Forskolin and Rolipram. 30 minutes after chLTP treatment, electrical activity was recorded for 5 minutes and a washout was done to remove the treatment: all medium was removed, followed by addition of 200 µl BrainPhys+SM1, all medium was removed again, finally 600 µl BrainPhys+SM1 media were added.

##### Immunocytochemistry

Dissociated organoids or stem cells were grown on round No. 1.5 coverslips placed in 12-well plates. The cells were cultured as normal and the cells were then fixed in 4% PFA with 4% sucrose in PBS with calcium and magnesium (+/+, Gibco 14040117) for 10 minutes at room temperature, washed twice in PBS (+/+), and then fixed in ice-cold methanol at 4 °C for 10 minutes. The cells were then washed 2x in PBS (+/+) and stored in PBS (+/+) at 4 °C until used. For staining, cells were permeabilised and blocked in 2% NGS or 2% NDS in PBS (+/+) with 0.1% Triton-X for 1.5 hours. Primary antibodies were diluted in the same buffer, but without Triton-X. Cells were incubated with primary antibodies at 4 °C overnight then washed 3x with PBS (+/+) for 15 minutes. The coverslips were then incubated with secondary antibodies diluted 1:1000 in 2% NGS/NDS for 1 hour at room temperature in the dark. The coverslips were then washed 3x with PBS (+/+) for 10 minutes and mounted onto microscope slides with Prolong Gold with DAPI as mounting media. The slides were allowed to cure overnight and were then sealed with nail polish.

##### Microscopy

Confocal microscopy was used to image organoid sections stained using IHC for regional and cell type identity markers and cells from dissociated organoids stained using ICC. Confocal images were acquired using an upright Nikon A1R confocal microscope with a 20x air objective, (N.A. 0.75) (www.kclwcic.co.uk/multiphoton). To image an entire section, a focus surface was set using the DAPI channel and the “scan large image” function was used to generate an assembled tile scan of one Z plane. Cells on coverslips were imaged as multiple regions of interest with a Z stack encompassing the cell layer. All acquisition parameters (including Z stack thickness, if applicable) were kept constant within an experiment.

#### RNA-seq

##### Cell lysis and RNA extraction

To extract total RNA from cells grown on the MEA from dissociated organoids, the medium was aspirated and 250 µl/well of accutase were added and the plate was incubated for 5 mins at 37 °C, 5% CO2. The cells with accutase from vehicle treatment condition were pipetted and pooled into a tube and an equal volume of HBSS was added. The cells were centrifuged at 1250 RPM and the medium was aspirated.

The cells were resuspended in 1 ml TRIzol^TM^, pipetted vigorously and transferred to a RNAse-free centrifuge tube. Each TRIzol^TM^ sample containing lysed cells was incubated at room temperature for five minutes. TRIzol^TM^ samples were stored at -80 °C until RNA could be extracted. To extract total RNA, samples were thawed at room temperature and incubated for 5 minutes at room temperature once thawed. 200 µl of chloroform was added to each sample, and the tubes were inverted and incubated for three minutes at room temperature. Finally, the sample was centrifuged at 10000 RPM for 5 mins at 4 °C and the aqueous phase was transferred to a new tube. 1 µl of glycogen and 500 µl isopropanol were added to the aqueous phase to increase RNA purity and yield. The sample was then inverted to mix, incubated for 15 minutes at room temperature, and centrifuged at 13000 RPM for 15 mins at 4 °C. The supernatant was discarded, and the pellet was washed by addition of 1 ml of 80% ethanol, centrifugation at 13000 RPM for 5 mins at 4 °C, and removal of the ethanol. Each pellet was air dried until transparent and resuspended in 30 µl RNAse-free H2O. To clean the RNA, the samples were then precipitated with 3M sodium acetate (Sigma S2889). 3 µl of sodium acetate and 90 µl 100% molecular-grade ethanol were added and the sample was incubated at -80 °C overnight. Finally, the sample was centrifuged at 13000 RPM for 15 mins at 4 °C and the supernatant was discarded. The pellet was washed with 80% ethanol two times, air dried until transparent, and then resuspended in 30 µl RNAse-free H2O. The RNA and protein concentration in each sample was measured using absorbance with a NanoDrop OneC spectrophotometer (Thermo Scientific ND-ONEC-W). The RNA was stored at -80 °C.

##### Bulk RNA sequencing

Purified RNA samples were sent for sequencing to Genewiz Inc (Azenta Life Sciences), who conducted additional quality control to check RNA integrity, size, and concentration using an RNA Qubit and Fragment Analyzer. Illumina sequencing libraries with PolyA selection were prepared in-house by Azenta. Sequencing was carried out using the Illumina NovaSeq in a 2x150bp configuration with an estimated data output of ∼20M paired-end reads per sample. Raw data was obtained from Azenta as FASTQ files.

### QUANTIFICATION AND STATISTICAL ANALYSIS

#### MEA data analysis

For data analysis, the .spk file was exported using Axis Navigator to the Neural Metrics Tool, where an analysis configuration was applied to extract individual MEA parameters (Table S1). The definitions of the MEA parameters (Table S1) are based on the definitions in the Axis Navigator User Guide, version 3.5^75^. The same analysis configuration was used across all recordings. To accommodate the lower network activity observed in organoids, we adjusted the default analysis configuration settings for bursting and network bursting. The analysis configuration used was as follows:

| CONFIGURATION | SETTING |
| --- | --- |
| Adaptive Threshold (SDs) | 5.5 |
| Active Electrode Criterion (spks/min) | 5 |
| Remove Coincident Artifacts | FALSE |
| *Electrode Burst Settings* |  |
| Algorithm | Poisson Surprise |
| Min Surprise | 3 |
| *Network Burst Settings* |  |
| Algorithm | ISI Threshold |
| Min # of Spikes | 35 |
| Max inter-spike interval (ISI) (ms) | 100 |
| Min # of Electrodes (%) | 25 |
| *Synchrony Settings* |  |
| Synchrony Window | 20 |
| *Average Network Burst Analysis Settings* |  |
| Network Burst Window Start (ms) | 0 |
| Network Burst Window End (ms) | 500 |
| Bin Size (ms) | 1 |

“Recommended metrics”, which include all standard parameters were exported to CSV files for statistical analysis in R (version 4.2.2). Raster plots of MEA activity with spike histograms were generated using Axis Navigator (Axion, version 3.9.1) and Neural Metrics Tool (Axion, version 4.0.5). Raw spike plots were generated using Axis Navigator. We used 6 and 37 DPD, along with three intermediate timepoints (15, 22, and 29 DPD), for more in-depth analysis of temporal changes in network activity. For the purposes of a consistent comparison of spontaneous activity, we focussed on four key parameters extracted from the Neural Metrics Tool across experiments, these being: *mean firing rate*, *number of bursts*, *area under normalised cross-correlation*, and *number of network bursts* (Table S1). These parameters each represent the primary parameter from each level of spontaneous activity (activity metrics, electrode burst metrics, network burst metrics, and synchrony metrics). Mean firing rate (the number of spikes over time (Hz)), was used as a general measure of spontaneous activity. Number of bursts was measured as the total number of single-electrode bursts across all electrodes in a well. Bursts were detected using a Poisson Surprise algorithm in the Axion Neural Metrics Tool, where neurons are assumed to fire according to a Poisson distribution, and the probability of a cluster of spikes appearing by chance is assessed using a “surprise” threshold ^76,77^. Area under normalised cross-correlation was used as a measure of synchrony across multiple electrodes. Area under normalised cross-correlation is the area under the curve of a pooled cross-correlogram across all unique combinations of electrodes in a well, normalised to auto cross-correlations (cross-correlation of an electrode to itself) to reduce effects of variability in firing at individual electrodes ^76,78^. Cross correlation measures the probability of a spike in one channel relative to a reference channel in a pair, with each spike being treated as time=0 in the reference, and surrounding spikes in the paired electrode being binned in a histogram. This parameter represents the likelihood of spikes happening together in pairs of electrodes and has been used as a readout of synchrony in previous organoid studies ^64^. As another measure of network activity, we measured the number of network bursts across all wells using an inter-spike interval (ISI) threshold method ^24,76^ with network bursts logged upon detecting at least 35 spikes across all electrodes in the well, each separated by an ISI of no more than 100 milliseconds with at least 25 percent of electrodes participating in the network burst. This parameter considers both the mean firing rate and the firing patterns across electrodes, providing an overview of the overall activity and synchrony of a neuronal network, and thus how coordinated or irregular the firing is across the network.

For functional network analysis, MEA network plots, and raster plots were plotted using MATLAB (version R2023b) and the MATLAB MEA network analysis pipeline (MEA-NAP, version 1.10.1)^26^. Waveform plots were generated using Axis Navigator. Analysis in MEA-NAP was done directly on .raw files exported from Axis Navigator (Axion). Raw files were first converted to .mat files using the file conversion tab in MEA-NAP. MEA-NAP was run using recommended settings, with sampling frequency set to 12500 Hz, potential difference units set to V, channel layout set to Axion 16, and wavelet cost of -0.12 ^79^. Spikes were detected using a template-based method, where a wavelet transform was used to identify spikes similar to a template wavelet (bior1.5 template). A spike time tiling lag (STTC) of 50 ms was used to determine correlation between spikes at different electrodes ^80^. A 50 ms STTC lag was used to take into account the larger spacing of electrodes, in order to detect network activity where not all neurons firing are detected. To remove correlations that occur by chance, the MEA-NAP pipeline detects significant functional connections using probabilistic thresholding. Network metrics exported by MEA-NAP were used for statistical analysis in R. We focussed on analysis of two basic node-level parameters extracted from MEA-NAP across experiments, these being mean node degree and mean edge weight (Table S1).

Unless otherwise stated, all MEA plots were generated using *ggplot2*^81^ in R from well average data exported from the Neural Metrics Tool or recording-level data for network activity exported from MEA-NAP.

#### Statistical analysis

Given that each well generated a unique neuronal network, each well was considered a biological replicate for the purpose of statistics, with batch and cell line, where applicable, used as covariates. Statistical analysis was performed using R version 4.2.2 in R Studio. All measured wells were included in the statistical analysis. Statistical analysis results are presented in Table S3, with significant effects being ones where p<0.05. All analysis for single-electrode parameters was performed on the well averages for all wells, e.g. the average value for all electrodes in a well.

For analysis of activity over time, the effect of timepoint was analyzed using a mixed-effects model with cell line as a fixed effect, followed by estimated-marginal means post-hoc comparisons for each cell line and DPD combination with Bonferroni correction for multiple comparisons (Table S3). This approach was used to account for the repeated measures design and fewer control wells at the 37 DPD timepoint.

For analysis of cell line differences at 37 DPD, analysis of variance (ANOVA) was done for each of six core parameters for the model *parameter ∼ line type (ESC or iPSC) + Cell line* or *parameter ∼ sex (XX or XY) + Cell line*. ANOVA tables are presented in Table S3, with significant effects being ones where p<0.05. Post hoc tests for pairwise comparisons between cell lines were performed using Tukey’s Honest Significant Difference method (Table S3).

For analysis of effects of chLTP or E2 (Table S3), we calculated the difference in the well average value of each parameter between baseline and 6 hours following chLTP, Vehicle, E2, or chLTP+E2 treatment, with N=12 per condition across two replicates from the same batch. Analysis of variance was done for each of six core parameters for the model *difference from baseline ∼ Condition + Replicate*. We selected this model over a model with interaction of replicate and condition using the Akaike Information Criterion (AIC). For parameters with a significant effect of condition, post hoc tests were performed using Tukey’s Honest Significant Difference method (Table S3). The same analysis method was used for effects of E2 compared to vehicle, where we compared the difference from baseline at 24 hours following the first treatment and 8 days following the first treatment (7 days of daily treatment) in the vehicle condition compared to E2. The 7-day E2/vehicle treatment experiment was performed in dissociated organoids from three cell lines (H1, H9, KOLF2.1J, and SCTi003-A), where N=6 per condition per line and batch.

Two batches were used for the KOLF2.1J and SCTi003-A lines.

For all linear regression analysis, diagnostic plots of residuals were plotted for each model to check assumptions of equal variance, outliers, normal distribution, and linearity. Data for some parameters in the temporal analysis, as noted in Table S3, was log-transformed as log(1+parameter) prior to statistical analysis.

#### Dimensionality reduction analysis

For dimensionality reduction analysis, we excluded wells without active electrodes (less than 5 spikes/minute). The following parameters were used for dimensionality reduction and clustering: network size, AUNCC, mean percentage of spikes in bursts, network density, global efficiency, ISI coefficient. of variation, inter-spike interval coefficient of variation, mean firing rate, mean node degree, number of network bursts, mean node strength, modularity score, mean significant edge weight. We selected parameters that could be measured across all wells and timepoints in active wells. Thus, we did not include parameters such as network burst duration, as such parameters are only measurable in wells with network bursting present. We also excluded parameters that were highly correlated. UMAP plots were generated using the umap R package^82^. Data for PCA analysis was normalized using the ‘scale()’ R function and principal components were calculated using the ‘princomp()’ function. Plots of variables were plotted using the factorextra^83^ package. Hierarchical clustering of the normalized parameters was performed using the ‘dist()’ R function to generate a matrix of euclidean distances, followed by clustering using ‘hclust()’ (average linkage method). PCA plots, dendrograms, and UMAPs were plotted using ggplot2.

#### Analysis of dissociated organoid culture morphology

To monitor morphology over time, a minimum of four wells per line and batch of dissociated organoids were imaged at 7, 13, 27, and 37 days after dissociation using an EVOS XL brightfield confocal microscope at 10x magnification. To quantify the ratio of sparse areas in the well with a cell monolayer to dense areas with cell clusters, a multi-class segmentation was applied to the brightfield images taken at 37 days post dissociation using the pixel classification function in Ilastik ^84^(1.4.0.post1), by training an Ilastik classification model on a subset of the images (11 training images). Areas of the well were segmented into dense areas, sparse areas, and electrodes. Segmented tiff format images were imported into FIJI 2.14.0/1.54F and the *glasbey* look-up table (LUT) was applied. Each image was converted to RGB, the channels split, inverted, converted to a mask, and the mask was used to create a selection, which was measured. A “cluster ratio” was calculated as the sparse area divided by the dense area. This ratio method uses the same principle as the method described by Hörberg *et al.* ^65^, but uses a different segmentation strategy. Segmented images for figures had the *glasbey inverted* LUT applied to be colourblind friendly.

#### Bulk RNA-seq analysis

The FASTQC tool (version 0.12.1) was used to do initial read quality control on the FASTQ files ^85^. FASTQ files were processed using command line tools on the King’s College London CREATE high-performance computing cluster. Alignment of reads to the human genome (Assembly GRCh38.p14, accession GCF_000001405.40, NCBI RefSeq) was performed using STAR (version 2.7.6a) ^86^. After alignment, the output BAM files were sorted by chromosome and position coordinates using Samtools (version 1.17) and Picard (version 2.26.2) was used to mark duplicates in the BAM files ^87,88^. The duplicate checked BAMs were downloaded from CREATE and subsequent analysis was carried out in R (version 4.2.2). A raw counts table was generated using the *featureCounts* function from the Rsubread package ^89^. Additional attributes were annotated using the *getBM* function from the *biomaRt* package, including Entrezgene ID, HGNC symbols, ENSEMBL gene IDs, gene size, chromosome and position ^90^. Differential gene expression analysis was performed using the *DESeq2* package ^91^. Heatmaps were generated using the *pheatmap* R package (version 1.0.12), dendrogram nodes were sorted based on subtree distance using the *dendsort* R package (version 0.3.4).

*Analysis of estimated organoid regional identity*

The *VoxHunt* ^63^package was used estimate the proportions of regional identities in each dissociated organoid sample. Gene lists representing brain regions were extracted from Allen Brain Atlas E13 *in situ* hybridization mouse data and were used to deconvolve the RNA-seq data using the *VoxHunt deconvolute* function. RNA-seq data was inputted as DESeq2 normalised counts for each sample.

## Notes

### Competing Interest Statement

The authors have declared no competing interest.

