## Supplemental Figures for "Electrophysiological development and functional plasticity in dissociated human cerebral organoids across multiple cell lines"

**Table S1. MEA parameter definitions**

| Parameter | Definition |
|---|---|
| **Spiking parameters** | |
| Mean Firing Rate (MFR)[a, b] | (Number of Spikes)/(Analysis duration) in Hz |
| Inter-spike interval (ISI) | The time between spikes. |
| ISI Coefficient of Variation[b] | (Standard deviation (SD) of ISI)/(mean ISI). Measure of spike regularity. |
| Network ISI Coefficient of Variation | Coefficient of variation (standard deviation/mean) of the ISI for all spikes on all electrodes in a well. Measure of spike regularity across the network, captures the distribution of spiking such that 0 indicates spikes perfectly distributed and >1 indicates network bursting. |
| Number of Active Electrodes | Number of electrodes with activity greater than the minimum spike rate of 5spks/min. |
| Weighted MFR | Mean firing rate using data from active electrodes only. |
| **Single electrode bursting parameters** | |
| Number of Bursts[a] | Total number of single-electrode bursts over a given duration. |
| Mean ISI within Burst | Average ISI, for spikes in a single electrode burst. Measure of burst intensity; smaller values mean more intense bursts. |
| Median ISI within Burst | Median ISI for spikes in a single electrode burst. |
| Inter-burst Interval | Average time between the start of single-electrode bursts. |
| IBI Coefficient of Variation | The coefficient of variation (standard deviation/mean) of the inter-burst interval, the time between single-electrode bursts. This is a measure of single electrode burst regularity. |
| Burst Percentage[b] | (Number of spikes in single-electrode bursts)/(total number of spikes)*100 |
| **Network bursting parameters** | |
| Number of Network Bursts[a,b] | Total number of network bursts over a given duration. |
| Network Burst Duration | Average time from the first spike to last spike in a network burst. |
| Number of Spikes per Network Burst | Average number of spikes in a network burst. |
| Number of Electrodes Participating in a Burst | Average number of electrodes with activity during a network burst. |
| Number of Spikes per Network Burst per Channel | (Average number of spikes per burst)/(number of electrodes participating in that burst) |
| Network Burst Percentage | (Number of spikes in network bursts)/(total number of spikes)*100 |
| Network IBI Coefficient of Variation | The coefficient of variation (standard deviation/average) for the inter-network burst interval. A measure of network burst regularity. |
| Network Normalized Duration IQR | Interquartile range of network burst durations. Provides a measure of network burst duration regularity. If the middle 50% of network bursts are |

| | |
|---|---|
| | approximately the same duration, this value will be small, whereas, if the network bursts vary widely in duration, this range will be large. |
| **Synchrony parameters** | |
| Area Under Normalized Cross-Correlation (AUNCC)[a, b] | Area under the well-wide pooled inter-electrode cross-correlation normalized to the autocorrelations. Higher areas indicate greater synchrony. |
| **Functional connectivity parameters** | |
| Mean Node Degree[a, b, c] | Mean number of significant connections (edges) of each node with other nodes (electrodes) in the network, determined using probabilistic thresholding. |
| Mean Edge Weight[a, b, c] | Mean Significant Edge Weight. Edge weight is the strength of connectivity between two nodes, calculated using a spike-time tiling coefficient of 50ms. |
| Modularity Score (Q)[b, c] | A value between -0.5 and 1 that describes how well a network has been partitioned. |
| Mean Node Strength (NSmean)[b, c] | Mean of node strength for all nodes in a well, where node strength is the sum of the edge weights for each node. |
| Global Efficiency (Eglob)[b, c] | Efficiency of parallel information transfer between nodes in the network. |
| Controllability[b, c] | Mean Average Controllability, metric of how much influence a node has over the overall network activity. |
| Network Density[b, c] | Number of connections (edges) as a proportion (%) of the total possible connections that can be formed in the network. |
| Network size (aN)[b, c] | Number of active electrodes in MEA-NAP pipeline |

Parameter definitions for parameters discussed in the study. Definitions for parameters calculated using the Axion Neural Metrics Tool are taken from the Axis Navigator User Guide, Axion BioSystems [1]. Definitions for parameters calculated using MEA-NAP are taken from Table 1 of Sit et al. [2].

[a]Core parameter used for analysis of activity and network characteristics
[b]Parameter included in main dimensionality reduction analyses
[c]Parameter calculated using MEA-NAP, Sit *et al.* [2]

**Table S2. Comparison of network parameters between two KOLF2.1J batches.**

| Parameter | Batch 1 (mean ± SD) | Batch 2 (mean ± SD) |
|---|---|---|
| Number of Network Bursts | 10.33 ± 3.08 | 2.83 ± 1.11 |
| Number of Spikes per Network Burst - Avg | 108.56 ± 43.19 | 686.42 ± 205.59 |
| Number of Elecs Participating in Burst - Avg | 7.69 ± 1.61 | 12.63 ± 2.09 |
| Number of Spikes per Network Burst per Channel - Avg | 13.99 ± 5.7 | 53.4 ± 13.3 |
| Mean ISI within Network Burst - Avg (sec) | 0.0158 ± 0.0041 | 0.0030 ± 0.0024 |
| Burst Percentage - Avg | 45.23 ± 8.46 | 72.29 ± 11.58 |
| Network Burst Duration - Avg (sec) | 1.29 ± 0.2 | 1.49 ± 0.29 |
| Median/Mean ISI within Network Burst - Avg | 0.66 ± 0.04 | 0.37 ± 0.07 |

Values are the mean of all well averages ± standard deviation at 30 DPD.

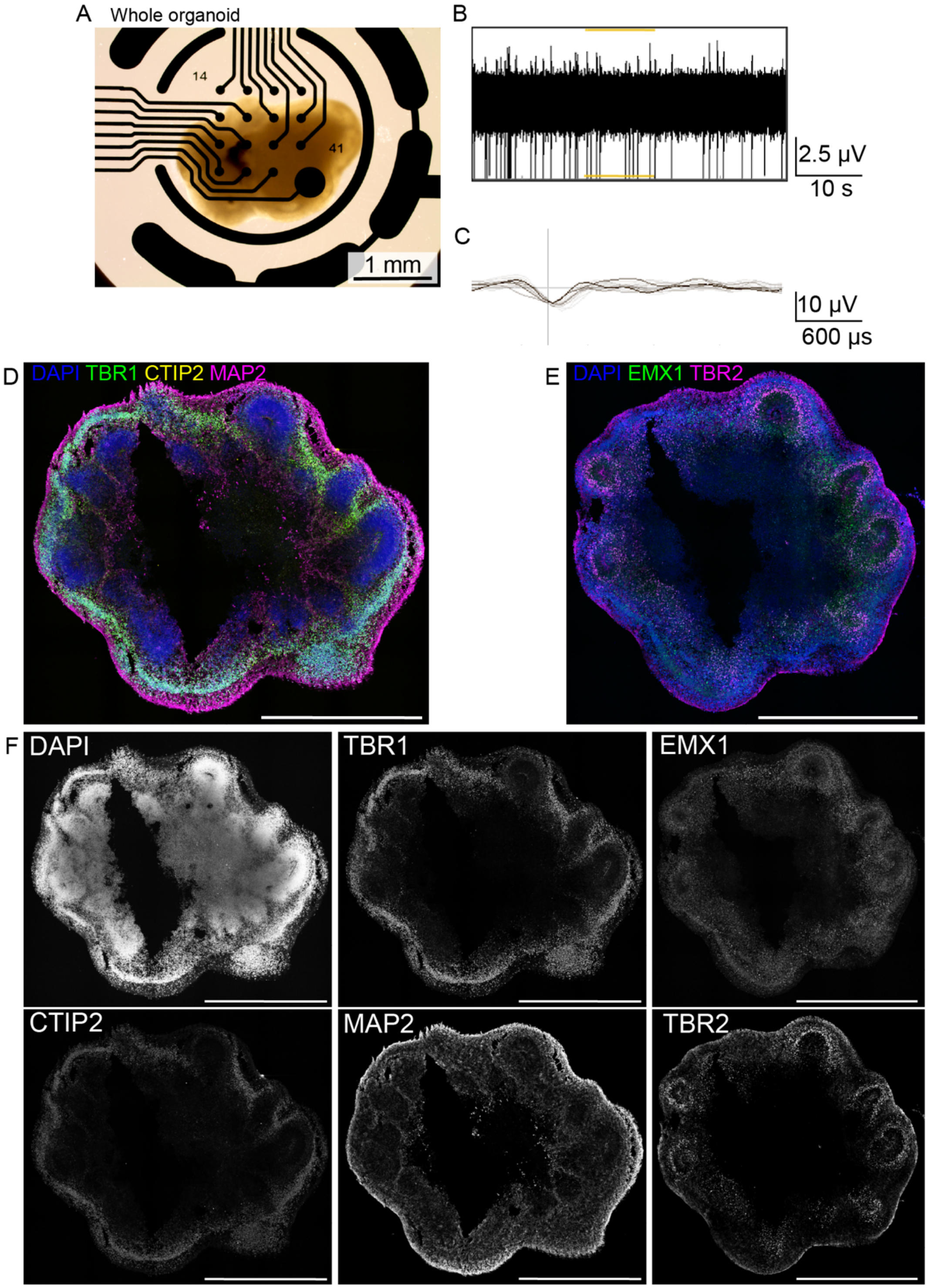


**Figure S1. IHC and MEA recording of cerebral organoids at 40 DIV. Related to Figure 1.**
(A) Brightfield image of whole cerebral organoid attached to a laminin-coated MEA plate.

(B-C) Snapshot of recording from H9 line whole organoid at 40 DIV showing unsorted overlaid spike waveforms recorded at a single electrode. Waveforms (C) correspond to individual spikes above the 5.5

standard deviation (SD) threshold (B). The darkest waveform represents the most recent spike recorded at the electrode. Waveforms were generated using Axis Navigator.

(D-F) Confocal tile scans of two adjacent serial sections of a representative SCTi003-A line organoid at 40 DIV. Immunostaining for dorsal forebrain markers EMX1 and TBR2 (E), MAP2 and dorsal forebrain markers CTIP2 and TBR1 (D). Corresponding grayscale images for each channel (F). Brightness and contrast were adjusted to allow markers to be distinguished. All scale bars represent 1 mm.

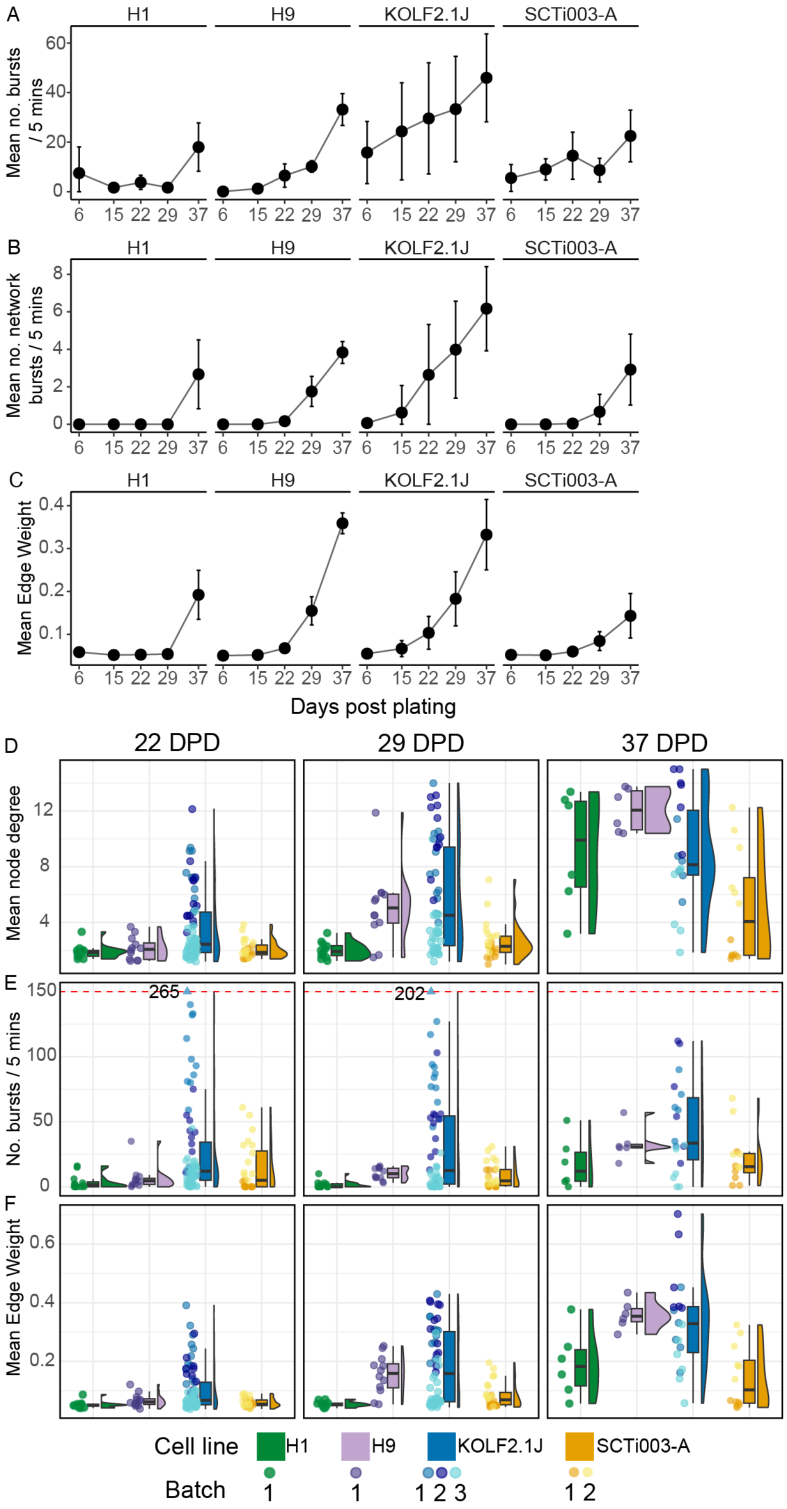


**Figure S2. Bursting and network connectivity increases over time, related to Figure 2.**
(A-C) Line graphs showing the mean number of bursts (A), mean number of network bursts (B), and mean edge weight (C) at 6-, 15-, 22-, 29-, and 37-days post dissociation (DPD). Each point represents the mean of well averages, the error bars represent the standard deviation of well averages. Full statistical analyses for these parameters are in table S3.

(D-F) Box and violin plots of the distribution, min, max, median, and interquartile range of mean node degree (D), mean number of bursts (E), and mean edge weight (F) per well at 22, 29, and 37 DPD. Each point represents the mean of one well across 5 minutes of recording. The colors of each point indicate the batch and cell line. The boxplot color indicates the cell line, ordered from left to right as H1, H9, KOLF2.1J, and SCTi003-A. For clarity, number of bursts (E) is capped at 150, two points with >150 bursts are shown above the dotted line and annotated with their actual values.

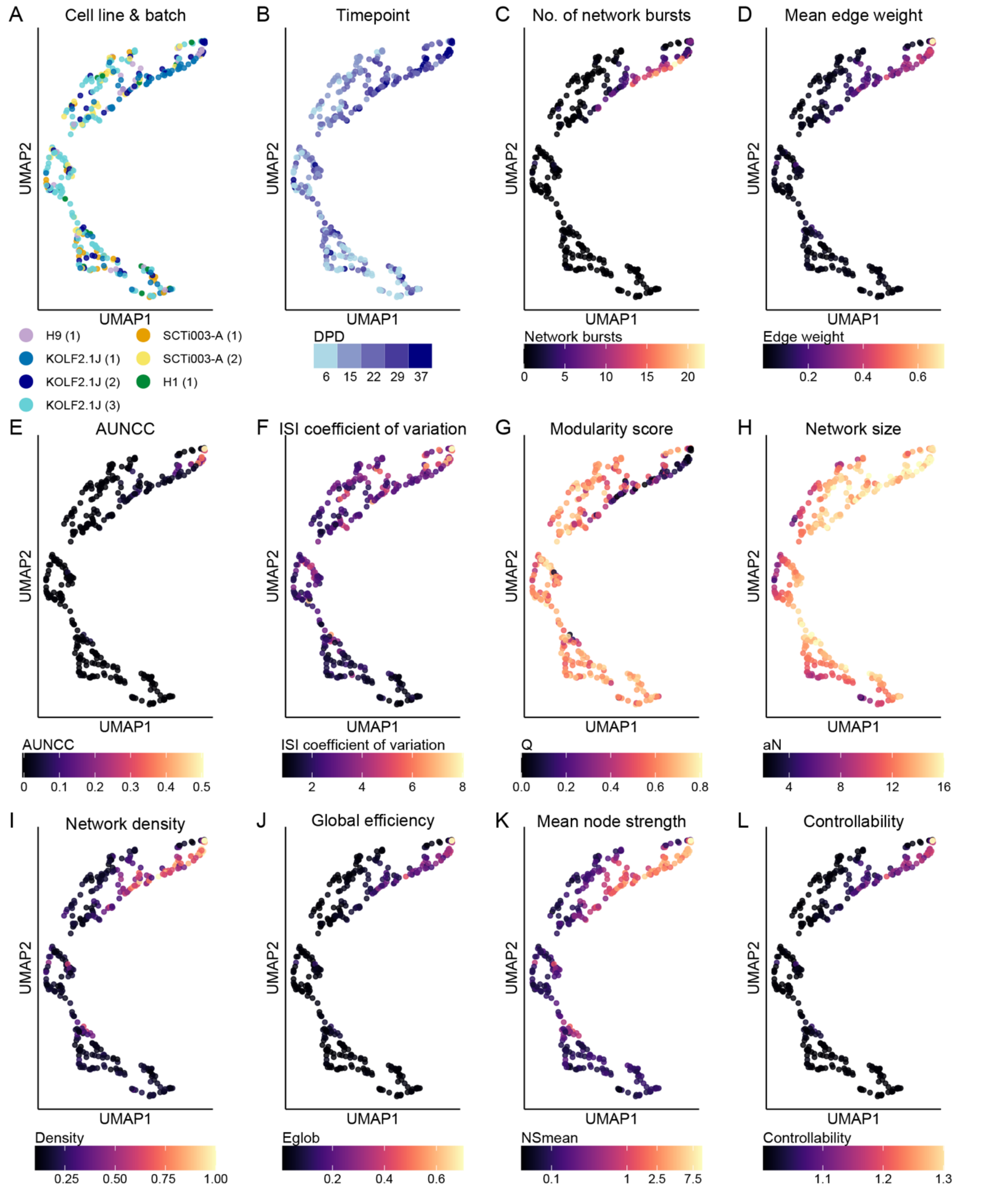


**Figure S3. UMAP visualizations of additional parameters included in dimensionality reduction analysis, Related to Figure 3**

(A-L) UMAP visualizations of MEA wells from 13 neuronal activity and network parameters showing all wells with active electrodes across five timepoints, 6-, 15-, 22-, 29-, and 37-DPD. Each point represents one well at one timepoint. Color-coded by timepoint (A), by cell line and batch (B), by network burst number (C), by mean edge weight (D), AUNCC (E), ISI coefficient of variation (F), modularity score (G), network size (H), network density (I), global efficiency (J), mean node strength (K) and controllability (L).

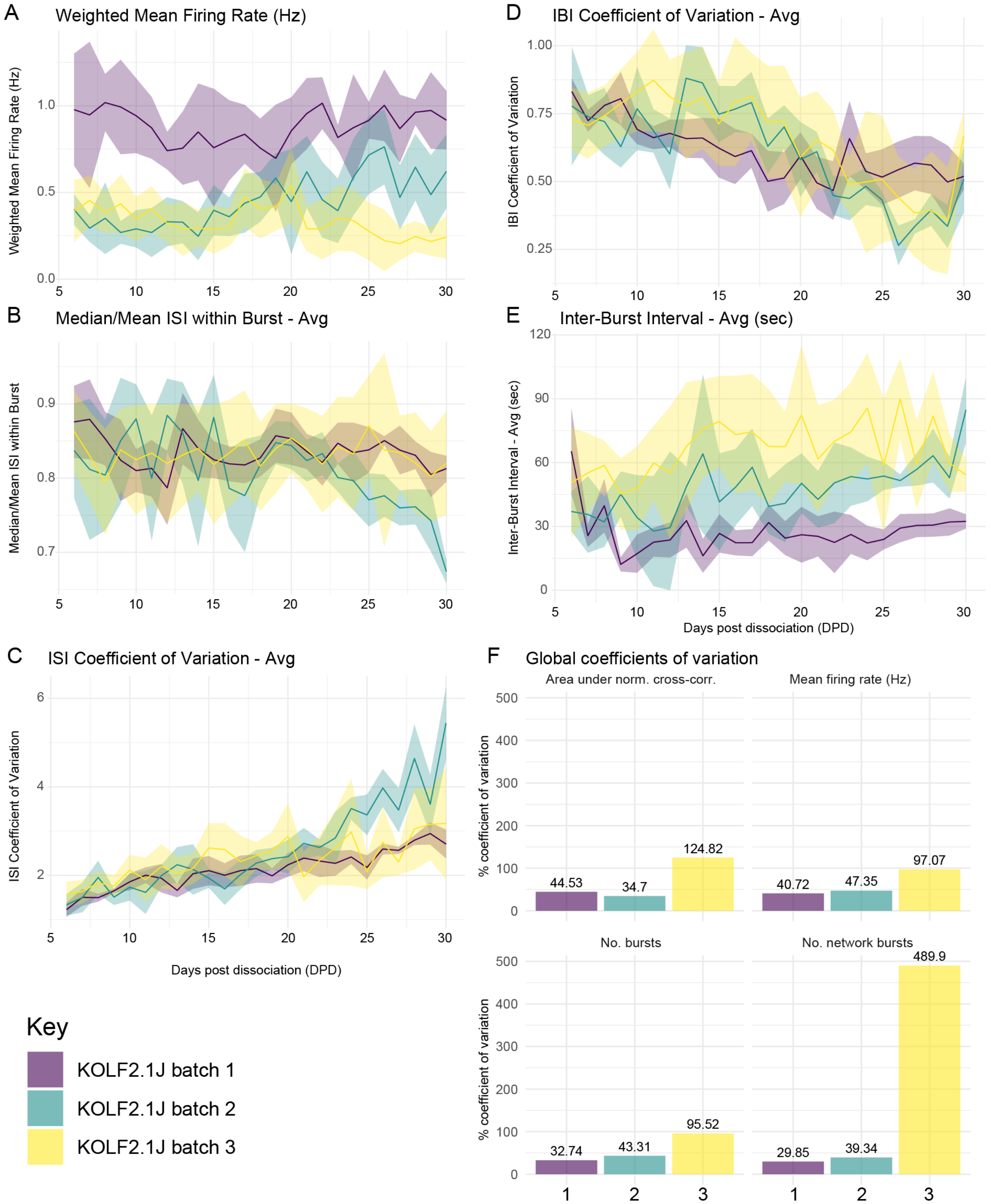


**Figure S4. Development of spontaneous electrical activity and network properties vary between batches – additional parameters and coefficients of variation, related to Figure 4**
(A-E) Line graphs of median across all wells for additional parameters over time showing the weighted mean firing rate (A), median/mean ISI within burst (B), ISI coefficient of variation (C), IBI coefficient of variation (D), IBI (E) across three independent batches of dissociated KOLF2.1J organoids, with one timepoint per day (5 mins recording). The ribbon represents the interquartile range.

(F) Bar graphs of “Global” coefficients of variation at 30 DPD for three batches of KOLF2.1J line dissociated organoids, for parameters shown in Figure 4. Coefficients of variation were calculated as the standard deviation of the well averages of each parameter divided by the mean of each parameter across all wells, multiplied by 100.

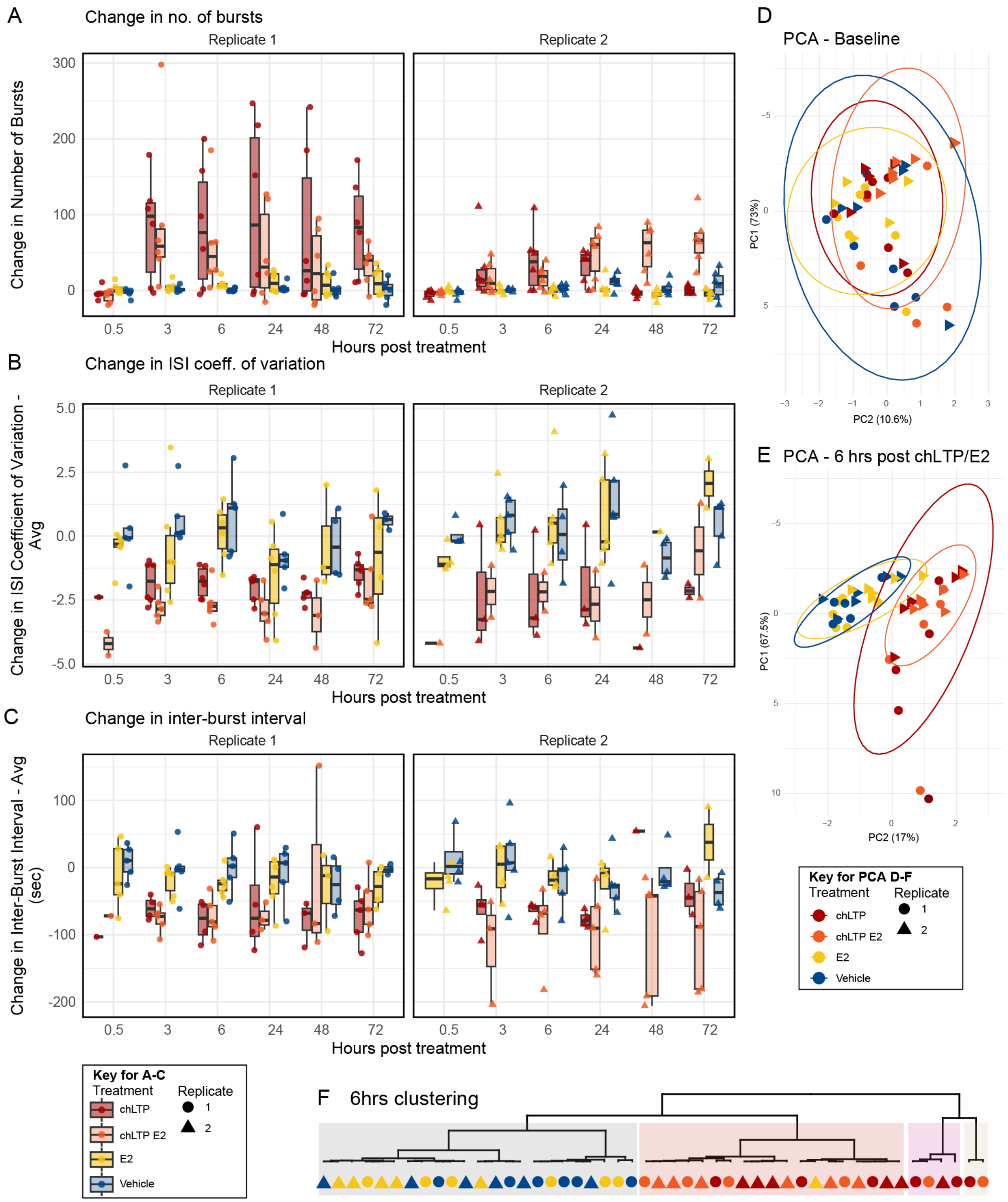


**Figure S5. Acute 17β-estradiol does not affect electrical activity in dissociated organoids in combination with a chemical plasticity stimulus, related to Figure 6**

(A-C) Boxplots of difference relative to baseline in MEA parameters after chLTP, chLTP+E2, E2, or vehicle treatment. Boxplots show the min, max, median, and interquartile range of the change in number of bursts / 5 minutes (A), ISI coefficient of variation (B), and inter-burst interval (C) per well, at six different timepoints (0.5, 3, 6, 24, 48, 72 hours post treatment. Each point represents the mean of one well across 5 minutes of recording. The color of each point indicates the treatment (vehicle green, chLTP red, chLTP+E2 blue, E2 yellow). For each parameter, the left plot shows the first replicate and the plot on the right shows the second replicate. In B-C, only wells with bursts present are shown. Measured in the KOLF2.1J, with treatment starting at 34 DPD (79 DIV) for replicate one and 29 DPD (74 DIV) for replicate 2. The data for chLTP and Vehicle is also shown in Figure 6.

(D-E) Unsupervised clustering analysis of MEA wells shows a clear effect of chLTP but no effect of E2 exposure. PCA analysis of 10 MEA parameters measured at baseline (A) and measured 6 hours following chLTP and E2 pre-treatment (B). The PCA plots show the first and second principal components, representing the two main sources of variation in the data. Each point represents the one MEA well, colored by treatment. Point shape indicates the replicate. The data ellipses represent the 95% confidence interval, expected to enclose 95% of bivariate-normal t distributed data for each treatment condition.

(F) Dendrogram of hierarchical clustering on principal components analysis for each well at 6 hours following chLTP shows vehicle/E2 and chLTP/chLTP+E2 samples clustering together. Clustering done using Ward’s criterion for the first three principal components.

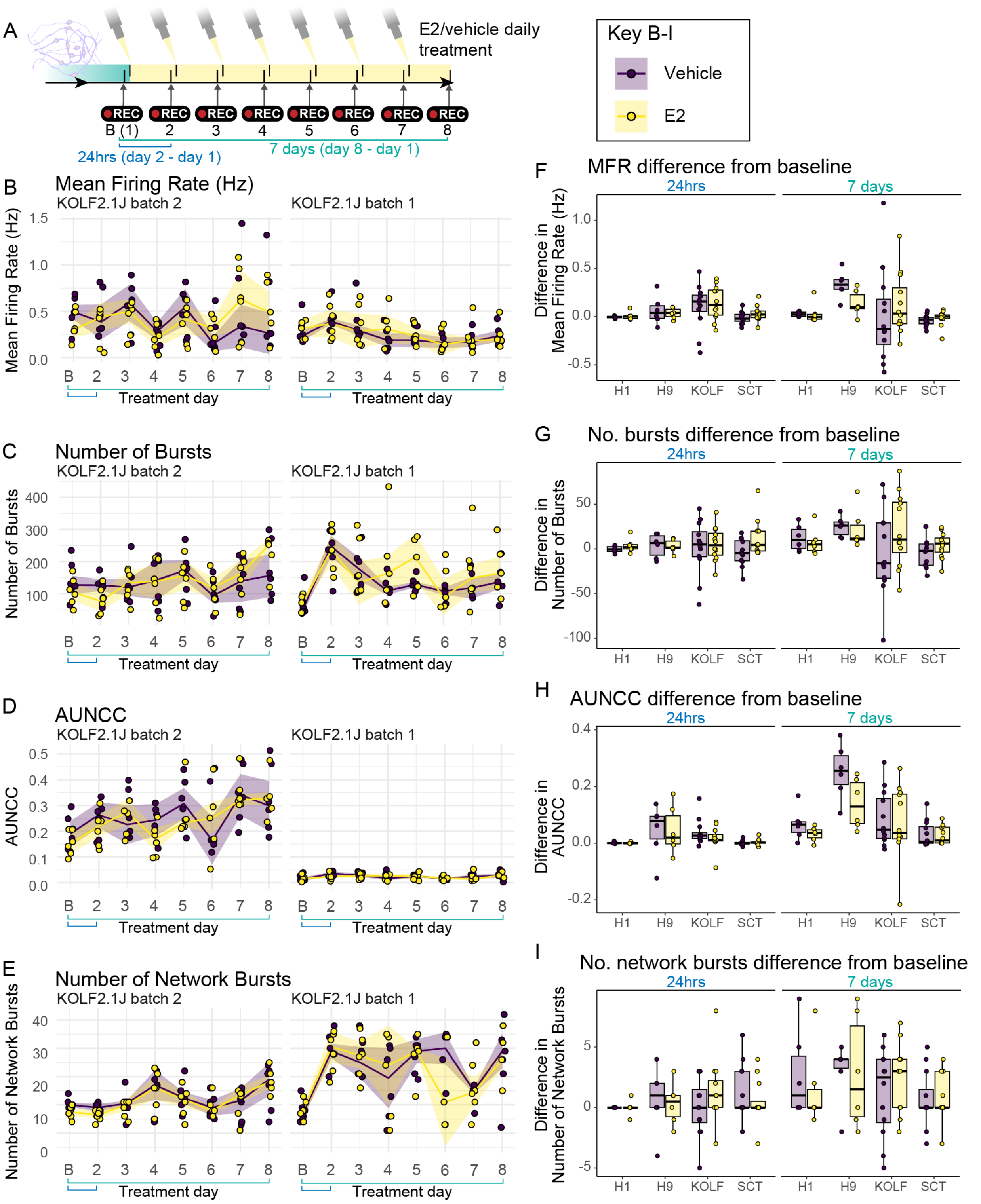


**Figure S6. Long-term 17β-estradiol exposure does not affect electrical activity in dissociated organoids, related to Figure 6**

(A) Schematic of treatment and recording timeline. The baseline recording was taken immediately prior to the first treatment on the first day. The next recording was taken 24 hours after the baseline, followed by another treatment spike. Recordings and treatments in subsequent days were performed in the same manner.

(B-E) Line graphs showing the median firing rate (B), number of bursts (C), area under normalized cross-correlation (D), and number of network bursts (E) across two independent batches of dissociated KOLF2.1J organoids, with one timepoint per day (5 mins recording). The ribbon represents the interquartile range. Each point represents one well. Wells without network bursting were excluded from the analysis. Point, line, and ribbon color indicates treatment (E2 yellow, vehicle purple). Treatment day 'B' is the baseline recording.

(F-I) Boxplots of the min, max, median, and interquartile range of the difference relative to baseline (day 1) in firing rate (F), number of bursts / 5 minutes (G), area under normalized cross- correlation (H), and number of network bursts / 5 minutes (I) per well after 24 hours or 7 days of daily treatment with E2/vehicle. Each point represents the difference in means of one well across 5 minutes of recording. The colors indicate the treatment. Experiment performed in dissociated organoids across two independent batches from the KOLF21.J and SCTi003-A line and one batch of the H1 and H9 line  starting at 29 DPD (baseline). Statistical analysis in Table S3.

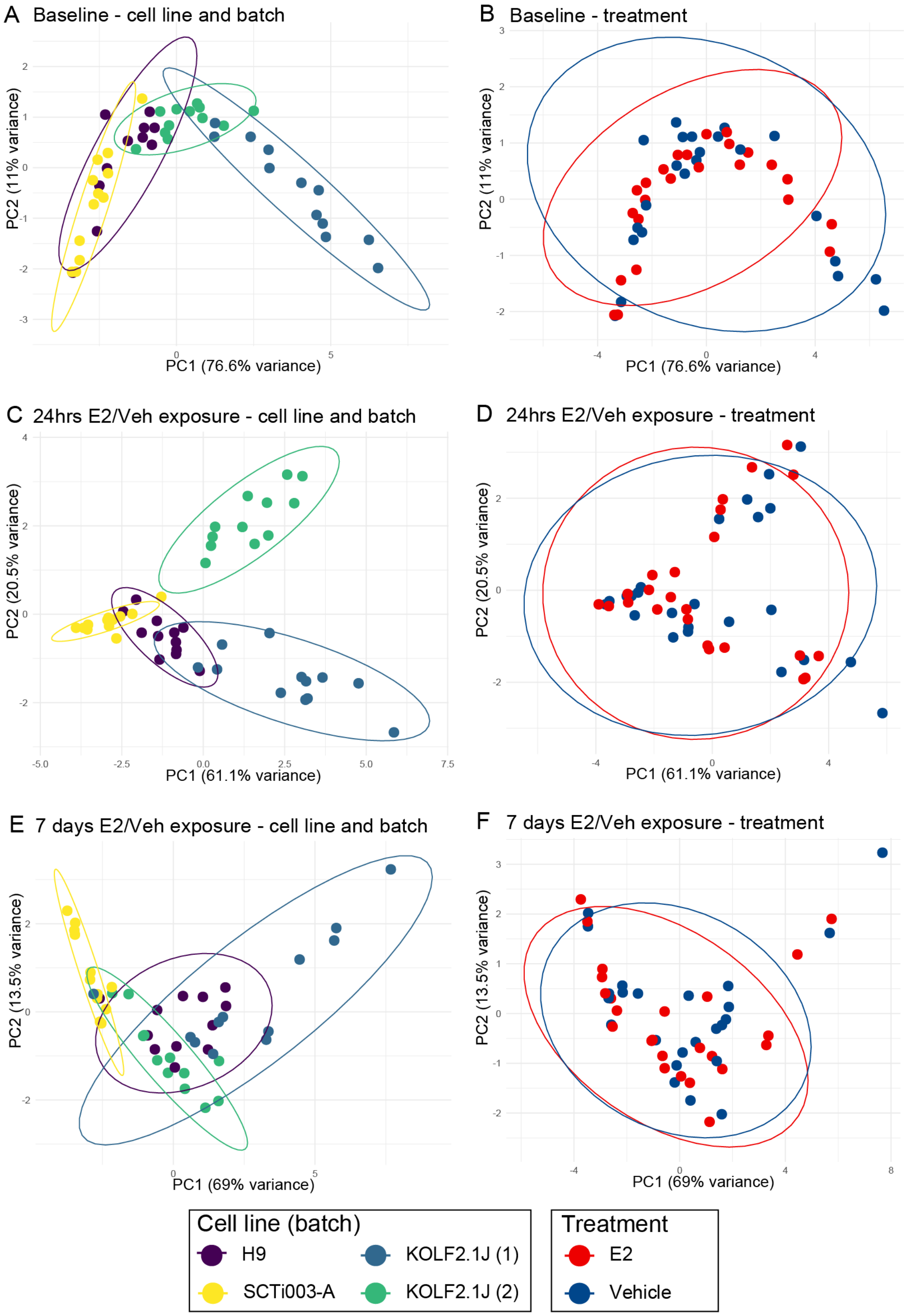


**Figure S7. Unsupervised clustering analysis of MEA wells shows batch and cell line differences but no effect of E2 exposure, related to Figure 6**

(A-F) PCA analysis of 10 MEA parameters measured at baseline (A-B), 24hrs E2 exposure (C-D), and 7 days E2 exposure (E-F). PCA plots of the first and second principal components, representing the two main sources of variation in the data. Each point represents one MEA well, colored by cell line and batch (left, A, C, E) and treatment (right, B, D, F). The data ellipses represent the 95% confidence interval, expected to enclose 95% of bivariate-normal t distributed data for each cell line/batch (left) or treatment condition (right). Experiment performed in dissociated organoids across two independent batches from the KOLF21.J line and one batch for the H1 and STi003-A line starting at 29 DPD (baseline), dissociated at 45-47 DIV.
